# Multi-Stable Bimodal Perceptual Coding within the Ventral Premotor Cortex

**DOI:** 10.1101/2024.12.11.628069

**Authors:** Bernardo Andrade-Ortega, Héctor Díaz, Lucas Bayones, Manuel Alvarez, Antonio Zainos, Natsuko Rivera-Yoshida, Alessio Franci, Ranulfo Romo, Román Rossi-Pool

## Abstract

Neurons in the primate ventral premotor cortex (VPC) respond to both tactile and acoustic stimuli, yet how they integrate and process information from these sensory modalities remains unclear. To investigate this, we recorded VPC neuronal activity in two trained monkeys performing a bimodal detection task (BDT), in which they reported the presence or absence of either a tactile or an acoustic stimulus. Single-cell analyses revealed diverse response types, including purely tactile, purely acoustic, bimodal, and neurons with sustained activity during the decision maintenance delay—the period between stimulus offset and motor response. To further examine VPC’s role in the BDT, we applied dimensionality reduction techniques to uncover low-dimensional latent dynamics in the neuronal population and conducted parallel analyses using a recurrent neural network (RNN) model trained on the same task. Neural trajectories for tactile and acoustic responses diverged sharply, whereas in stimulus-absent trials, the dynamics remained at rest. During the delay period, the trajectories exhibited a pronounced rotational dynamic, shifting toward a subspace orthogonal to the sensory response space, suggesting memory maintenance through stable equilibria. This indicates that network dynamics can sustain distinct stable states corresponding to the three potential task outcomes. Using low-dimensional modeling, we propose a universal dynamical mechanism that underlies the transition from sensory processing to memory retention, aligning with both experimental and computational findings. These results demonstrate that VPC neurons encode bimodal information, integrate competing sensory inputs, and maintain decisions across the delay period, regardless of sensory modality.

## INTRODUCTION

In a noisy restaurant, you can still hold a conversation with friends despite the ambient clamor. The brain effectively tunes into one speaker and filters out background noise– partly by integrating auditory signals with visual cues like lip movements (*1*). This natural interplay between sensory modalities demonstrates how the brain routinely combines information from different senses. While some cortical areas are specialized for multimodal integration (*1*), others resolve competition among heterogeneous sensory inputs. Understanding how the brain processes and integrates such information remains a vibrant area of research.

Historically, multisensory processing was thought to rely on extensive unimodal processing within separate sensory cortices (*2, 3*). However, recent studies suggest that even primary sensory areas–once believed to handle only a single type of input– contribute to multisensory integration (*4–6*). Evidence for this comes from investigations using various measures of neural activity, ranging from low spatial resolution (EEG, fMRI) (*7–9*) and medium resolution (LFP) (*5, 10*) to high resolution (single-unit recordings) (*5, 11, 12*). Although multisensory processing can occur in early cortical regions, the full encoding of perceptual events appears to emerge in higher-order areas within the processing hierarchy (*13–16*).

In our previous work, we addressed multimodal sensory integration by recording single-unit activity in non-human primates performing a bimodal detection task (BDT) to determine whether areas 3b and 1 of the primary somatosensory cortex (S1) exhibit multisensory processing (*17*). In this task (Fig. 1A), monkeys were required to report the presence or absence of a sensory stimulus–which could be either tactile or acoustic–and if detected, to indicate its modality. Tactile and acoustic stimuli of varying amplitudes were randomly interleaved with stimulus-absent trials so that each trial presented either no stimulus or a stimulus (tactile or acoustic) at above-threshold, near-threshold, or sub-threshold levels (*18*). Thus, the BDT compels the brain to weigh tactile and acoustic signals against each other to generate the correct response, requiring the integration of both modalities. Although we identified hierarchical differences between areas 3b and 1, both predominantly exhibited pure tactile responses. This observation raised the question of which cortical area might encode and integrate both tactile and acoustic inputs.

**Figure 1.**
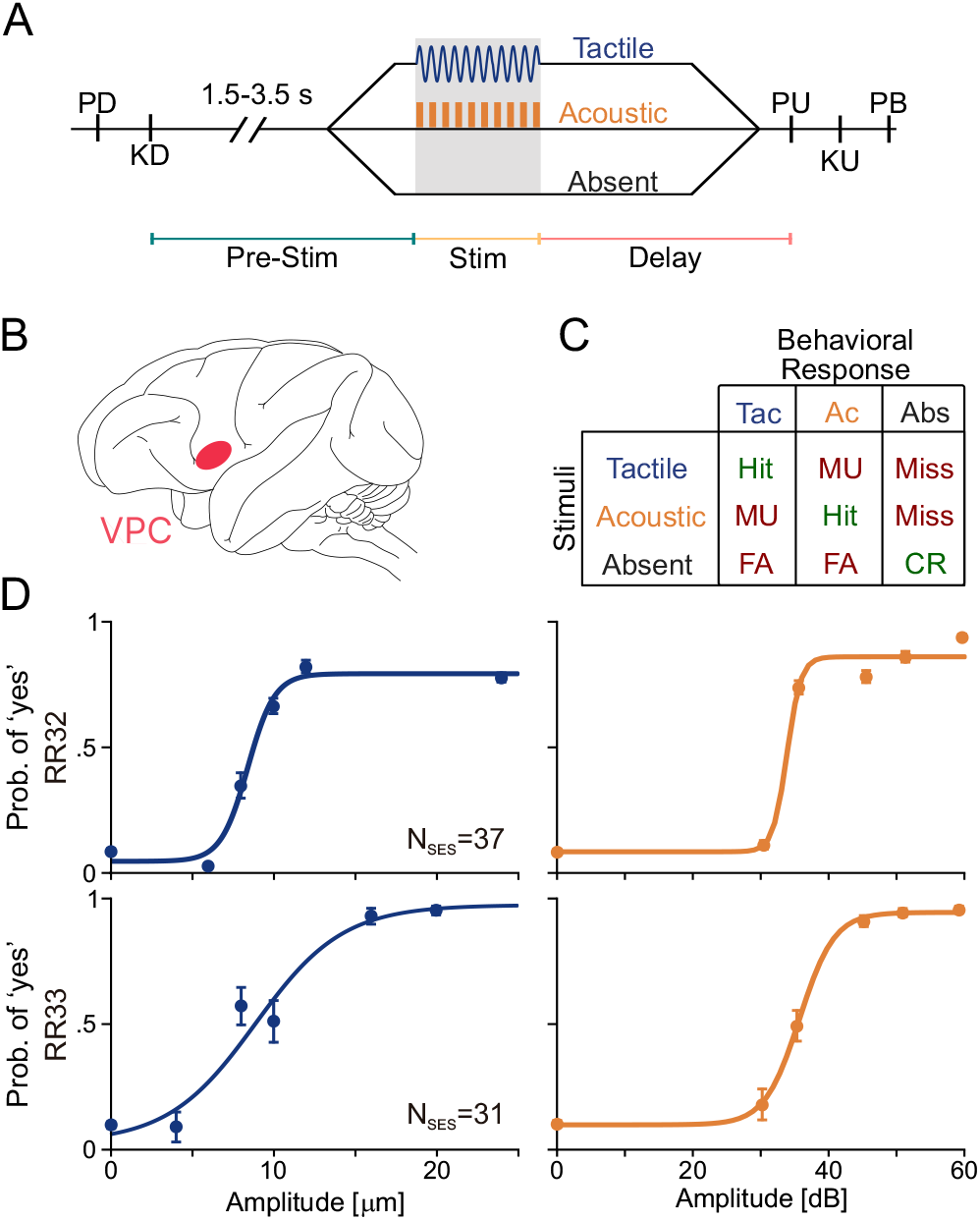
(A) Schematic of the Bimodal Detection Task (BDT). Trials began with the lowering of a mechanical probe and the indentation of the glabrous skin on one fingertip of the monkey’s restrained right hand (Probe Down event, “PD”). The monkeys then placed their left free hand on a non-moving key (Key Down event, “KD”). After KD, a variable period (1.5 to 3.5 s) ensued to prevent anticipation of the upcoming stimulus. This was followed by a 0.5 s stimulus period, which could be tactile, acoustic (20 Hz), or absent. Post-stimulation or stimulus-absent period, a fixed 2 s delay occurred, leading to the probe’s retraction (Probe Up event, “PU”), signaling the monkey to release the key (Key Up event, “KU”) and report its decision by pressing one of three buttons placed at eye level in front of them (Push Button event, “PB”), indicating “tactile stimulus present,” “acoustic stimulus present,” and “stimulus absent.” Correct responses were rewarded with a few drops of fruit juice. (B) Depiction of the left cerebral hemisphere with the VPC delineated in red. (C) Matrix categorizing trial outcomes by stimuli and behavioral responses. Correct identifications are denoted in green, while incorrect responses are colored red. In the absence of a stimulus, an incorrect “stimulus present” response indicates a false alarm (FA), while a “stimulus absent” response results in a correct rejection (CR). With a stimulus present, a “present” response that aligns with the stimulus modality constitutes a hit. A “present” response that does not match the modality is a misunderstood trial (MU), and an “absent” response is a miss. (D) Psychometric curves for tactile (in blue) and acoustic (in orange) stimuli. Each modality features six classes of stimulus intensity from 0 to 24 μm (tactile) and 0 to 59 dB (acoustic), over 37 sessions for subject RR32 and 31 sessions for subject RR33.

Previous research has identified the VPC as a potential intermediary hub between sensory and frontal cortices (*2, 19*). The VPC receives projections from sensory regions in the parietal and temporal lobes, as well as from association areas in the prefrontal and premotor cortices (*20–23*). In naïve monkeys, some VPC neurons respond to both tactile and acoustic stimuli (*24, 25*). During tactile tasks, VPC units display sensory, categorical, and persistent decision-related responses (*3, 19, 26*), and similar response patterns are observed during acoustic tasks (*27*). These findings suggest that the VPC is well-equipped to process and relay multimodal sensory information. Yet, it remains unclear whether– and how–the VPC contributes to the comparison of competing sensory modalities as required by the BDT. This uncertainty raises several key questions: Do VPC neurons encode both sensory modalities? How diverse are their response patterns? And can we distinguish between population-level responses to different types of stimuli?

In this study, we recorded neuronal activity from the VPC of two monkeys performing the BDT, analyzing the data at both single-neuron and population levels. We identified unimodal neurons that encoded the presence of either acoustic or tactile stimuli, as well as bimodal neurons that signaled stimulus presence for both modalities. Notably, the responses of bimodal neurons could also be interpreted as amodal, potentially aiding in the detection of stimulus absence. By applying dimensionality reduction techniques, we uncovered low-dimensional neural dynamics in the VPC that clearly differentiated responses to each sensory modality during stimulation. Specifically, acoustic and tactile stimuli elicited sharply diverging trajectories within a sensory subspace, while the population dynamics remained stable during stimulus-absent trials. Later, decision-related dynamics emerged orthogonal (uncorrelated) to the sensory responses, exhibiting a rotational pattern toward a memory subspace–presumably to reduce interference between sensory and mnemonic processing (*28, 29*).

Furthermore, a recurrent neural network (RNN) (*30, 31*) model trained on the BDT exhibited similar low-dimensional neural dynamics. This finding suggests that the observed diverge-then-rotate dynamics may represent a universal computational strategy for sensory discrimination followed by memory retention with minimal sensory-mnemonic interference. Through low-dimensional dynamical system modeling, we show that these dynamics arise from two saddle-point thresholds–each linked to one stimulus modality–and the geometry of their stable manifolds within a simple, multi-stable feedforward circuit. Together, our results implicate the VPC as a critical hub for integrating sensory inputs across modalities to guide perceptual decisions.

## Results

Two rhesus monkeys (Macaca mulatta) were trained to perform BDT in which they determined the presence or absence of sensory stimuli–either vibrotactile (Tac) or acoustic (Ac)–ranging from barely detectable to clearly perceptible. Vibrotactile stimuli were delivered at 20 Hz for 0.5 seconds (see Fig. 1A), whereas acoustic stimuli were 1 kHz pure tones modulated at 20 Hz, also lasting 0.5 seconds. In tactile trials, the amplitude of the skin vibrations were precisely controlled; similarly, the intensity of the tones was carefully regulated during acoustic tests (*17*). Sessions alternated between these sensory modalities and included an equal number of no-stimulus (Abs) trials. Following a decision-maintenance period, a “probe up” signal (“pu”) indicated that the animal could raise its hand (“ku”) and press the decision button (“pb”). The monkeys signaled their detection (or non-detection) of the stimulus by pressing one of three push buttons labeled “Tac”, “Ac”, or “Abs”, which were conveniently positioned along their left midline. VPC’s neuronal activity was recorded throughout the sessions (Fig. 1B). Trials were meticulously classified based on stimulus type and the corresponding behavioral response (Fig. 1C), and psychometric analysis confirmed that the monkeys consistently performed near their psychophysical thresholds (Fig. 1D).

### Individual neuron responses

We recorded neural activity from the VPC in animals RR32 (n=256 from the right hemisphere and n=310 from the left hemisphere) and RR33 (n=345 from the left hemisphere), yielding a total of 911 units. In contrast to our previous findings in areas 3b and 1 of the S1–where neurons exhibited highly faithful responses exclusively to tactile stimuli (*17*)–the VPC population displayed transformed responses to both modalities, indicating a distinct integration of acoustic and tactile inputs (Figs. 2 and S1). Some units showed selective responses to a single sensory modality, responding exclusively to either tactile or acoustic stimuli. For example, the unit depicted in Fig. 2A demonstrates a strong preference for acoustic stimuli, with its response remaining independent stimulus amplitude. This characteristic suggests that the neuron encodes the presence of acoustic stimuli rather than variations in intensity or other stimulus attributes. Similarly, the neurons illustrated in Figures 2B & D and S1B responded selectively to tactile stimuli. Conversely, units such as the one shown in Fig. 2C specifically encoded stimulus absence by exhibiting a reduced firing rate during both tactile and acoustic trials. However, most units displayed a variety of response patterns that varied with stimulus type, as exemplified by the neurons in Figs. 2E and S1G. Additionally, some neurons (e.g., those shown in Figs. 2F and S1I) exhibited decision-making-related responses during the movement period following the “pu” signal. In summary, the VPC exhibits a wide variety of responses patterns, with the majority of the units primarily encoding the presence of tactile or acoustic stimuli, both modalities simultaneously, or the stimulus absence.

**Figure 2.**
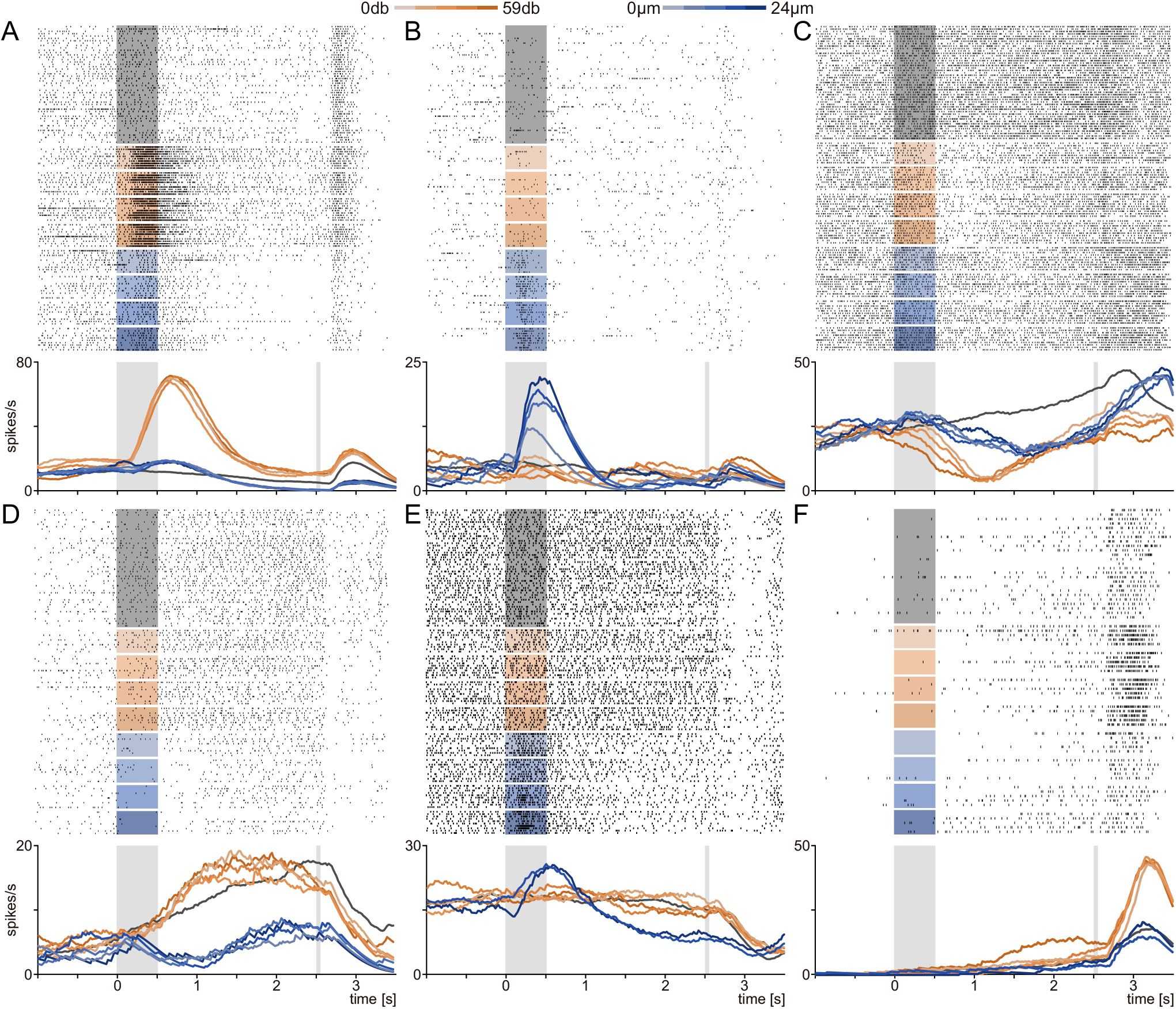
(A–F) Raster plots and activity profiles of six representative neurons recorded in the VPC. Tactile and acoustic trials are represented by shades of blue and orange, respectively, according to stimulus amplitude, while a deep-gray shade indicates stimulus-absent trials. Black ticks represent neuronal spikes, and light-gray rectangles indicate the stimulation period and PU event. Colored traces depict averaged neuronal activities aligned within trials sharing identical stimulus classes, which enables comparisons across different stimulus intensities and sensory modalities.

### Single unit cognitive dynamics

To obtain a population-level overview of these heterogeneous responses, we first computed the variance across different amplitude levels for tactile (*Var*_*Tac*_) and acoustic (*Var*_*Acu*_) stimuli (Fig. 3A, blue and orange traces, respectively), as well as the variance across the three decision outcomes (*Var*_*Dec*_; Fig. 3A, purple trace; see Methods). After subtracting the baseline variance (from t = -2 to -1 s), we found that all metrics peaked shortly after stimulus presentation–with a notably larger increase for the tactile condition– and remained stable during the delay until the “pu” event, after which all metrics rose again, likely reflecting decision execution. The condition-independent variance (*Var*_*Temp*_; Fig. 3B, green trace) followed a similar temporal profile but was consistently higher than the condition-specific variances, suggesting that most variability captured the sequential task events rather than differences modality or amplitude conditions. Likewise, the analysis of the stimulus-related information (Fig. 3C) showed that coding began shortly after stimulus presentation, persisted throughout the delay, and increased again after “pu”, highlighting significant VPC population coding throughout the task.

**Figure 3.**
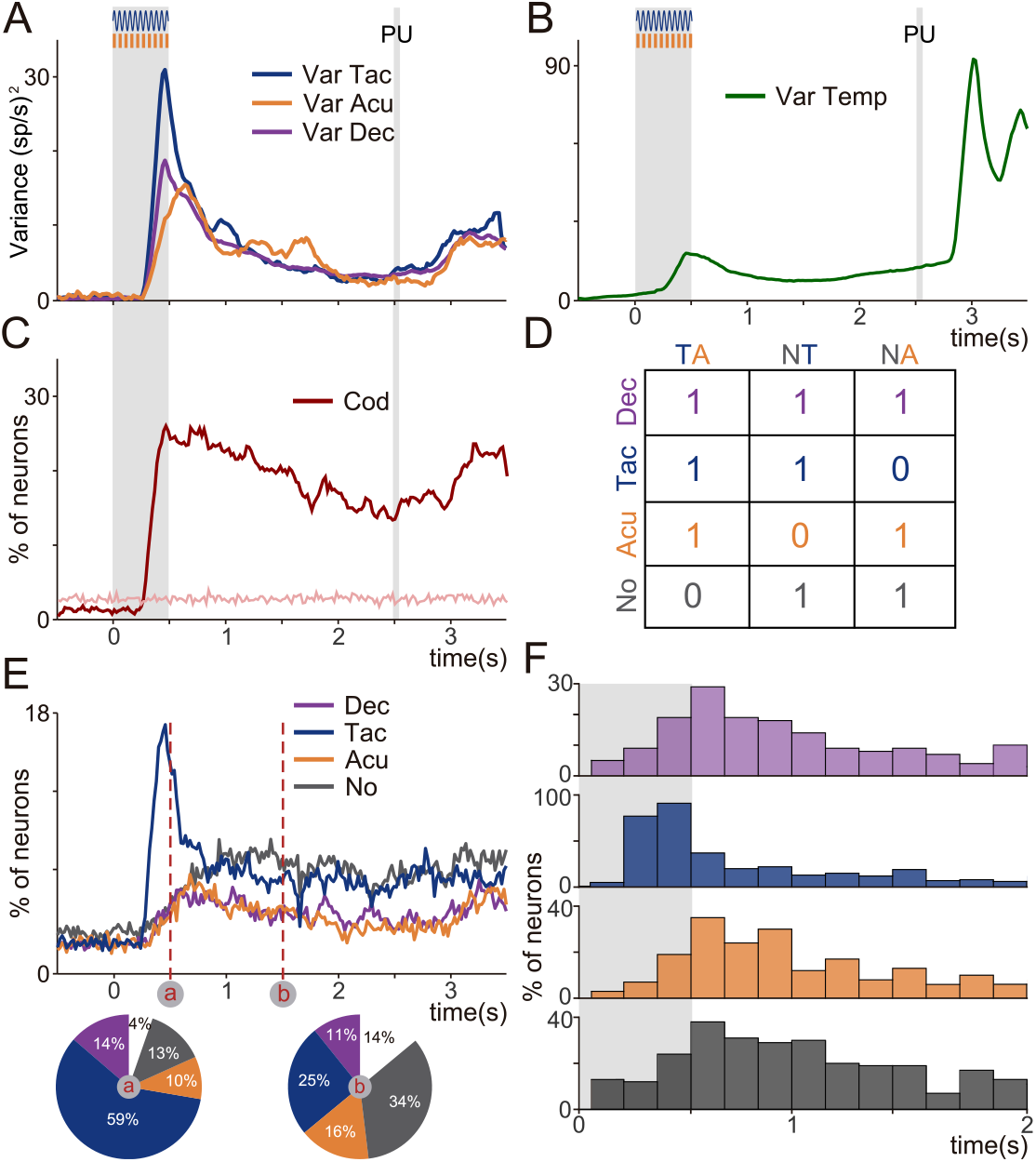
(A) Cognitive population variances as a function of time for the average response to each tactile (*Var*_*Tac*_, blue trace) and acoustic (*Var*_*Acu*_, orange trace) stimulus class, as well as for the mean response to each decision type (*Var*_*Dec*_, purple trace). The grey-shaded rectangles denote the periods of stimulation and the PU event. Basal variances (associated with residual fluctuations) were subtracted in all computed variances. (B) Temporal (or non-coding) variance (*Var*_*Temp*_, green trace), capturing the sequential events of the task rather than variations across classes within modalities or decisions. (C) The percentage of neurons conveying significant information about distinct decision outcomes. For this assessment, only units demonstrating significant information were included. Comparisons evaluated were: Tactile (T) versus Acoustic (A), Tactile (T) versus Absent (N), and Acoustic (A) versus Absent (N). The pink trace represents the baseline variability of neural activity observed across the population. (D) Matrix categorizing neurons by their type of encoded information within the indicated comparisons TA, NT, and NA. For each comparison, each neuron received a ‘1’ for significant information encoding or a ‘0’ for nonsignificant. Units were designated as ‘decisional’ (Dec) for scoring ‘1’ in all comparisons, ‘tactile’ (Tac) for ‘1’s in TA and NT, ‘acoustic’ (Acu) for ‘1’s in TA and NA, and ‘absent’ (No) when ‘1’s were exclusive to NA and NT. ((E) Percentage of neurons carrying significant information according to the classification made in (D). (F) Upon the presentation of a stimulus, most neurons predominantly encode tactile information. This is evident in the caked plot ‘a’, calculated using the final two time-bins of the stimulation period. As the delay epoch progresses, the unit’s information encoding becomes more uniform among the four encoding types Dec, Tac, Acu, No (‘b’ pie chart computed at 1.5s). The white area represents neurons with significant coding that could not be classified within (D). (G) Coding latency for different labeled subgroups of neurons (Dec, Tac, Acu, or No). Consistently with the analysis in (E), neurons carrying significant tactile information displayed faster encoding compared to other neurons.

To further examine how VPC neurons represented task-related information, we segmented neural responses into 200-ms bins and classified each response into one of four groups–decisional (Dec), tactile (Tac), acoustic (Acu), and stimulus absent (No)– based on significant mutual information comparisons (Fig. 3D: TA for tactile-acoustic; NT for absent-tactile; NA for absent-acoustic). For instance, neurons with significant differences in all comparisons were deemed decisional, while those significant only for TA and NT were categorized as tactile, and so on. This method revealed that tactile encoding was the most dominant and emerged earliest (Fig. 3E, pie chart “a”), though during the delay, coding became evenly distributed across categories (Fig. 3E, pie chart “b”). Additionally, coding latency analysis (Fig. 3F) showed that tactile neurons responded earlier than neurons encoding other information, underscoring VPC’s integrative capacity.

Finally, splitting the data by hemispheres and subject (Fig. S2A-C) confirmed that similar coding signals were present across all groups–even when tactile processing occurred in one hemisphere and motor commands in the other. Overall, these findings indicate that the VPC initially exhibits a strong tactile response, then shifts to a bimodal coding regime during the delay where neurons encode tactile, acoustic, stimulus absent, or decisional information. This dynamic coding reflects the VPC’s flexible role in integrating sensory inputs to guide perceptual decisions.

### Population dynamics during BDT

To elucidate the global interactions underlying the diverse VPC responses, we applied dimensionality reduction to reveal the latent dynamics of multisensory integration. We represented the VPC population as a point in an n-dimensional space (n=911 neurons) by combining firing rates from different experiments into a population vector (*29, 32–34*). Although this space is high-dimensional, the neural activity is assumed to evolve near a low-dimensional neural manifold that provides insight into population-level computation (*35–38*). Here, we used several methods to identify, visualize, and interpret the structure of the VPC neural manifold during the BDT.

We first employed principal component analysis (PCA) to characterize population dynamics over the entire task (Fig. 4A), the sensory (stimulus arrival) phase (Fig. 4B), and the memory (delay) (Fig. 4C). For each phase, covariance matrices were computed (see Methods), and significant PCs were identified by comparing their variance to that from a noise covariance matrix (p<0.01) (*33, 39*). Analysis across the entire task revealed that only the first seven PCs were significant (Fig. 4D, top). Projecting activity onto the first six PCs (Fig. 4A) showed that the 1^st^ PC primarily captured tactile responses, the 2^nd^ PC mainly reflected acoustic (though partially bimodal) responses, and later PCs were linked to decision-related signals during the delay. Notably, the 3^rd^ PC alone could decode the absence of stimuli, while 4^th^ distinguished among the three possible decisions.

**Figure 4.**
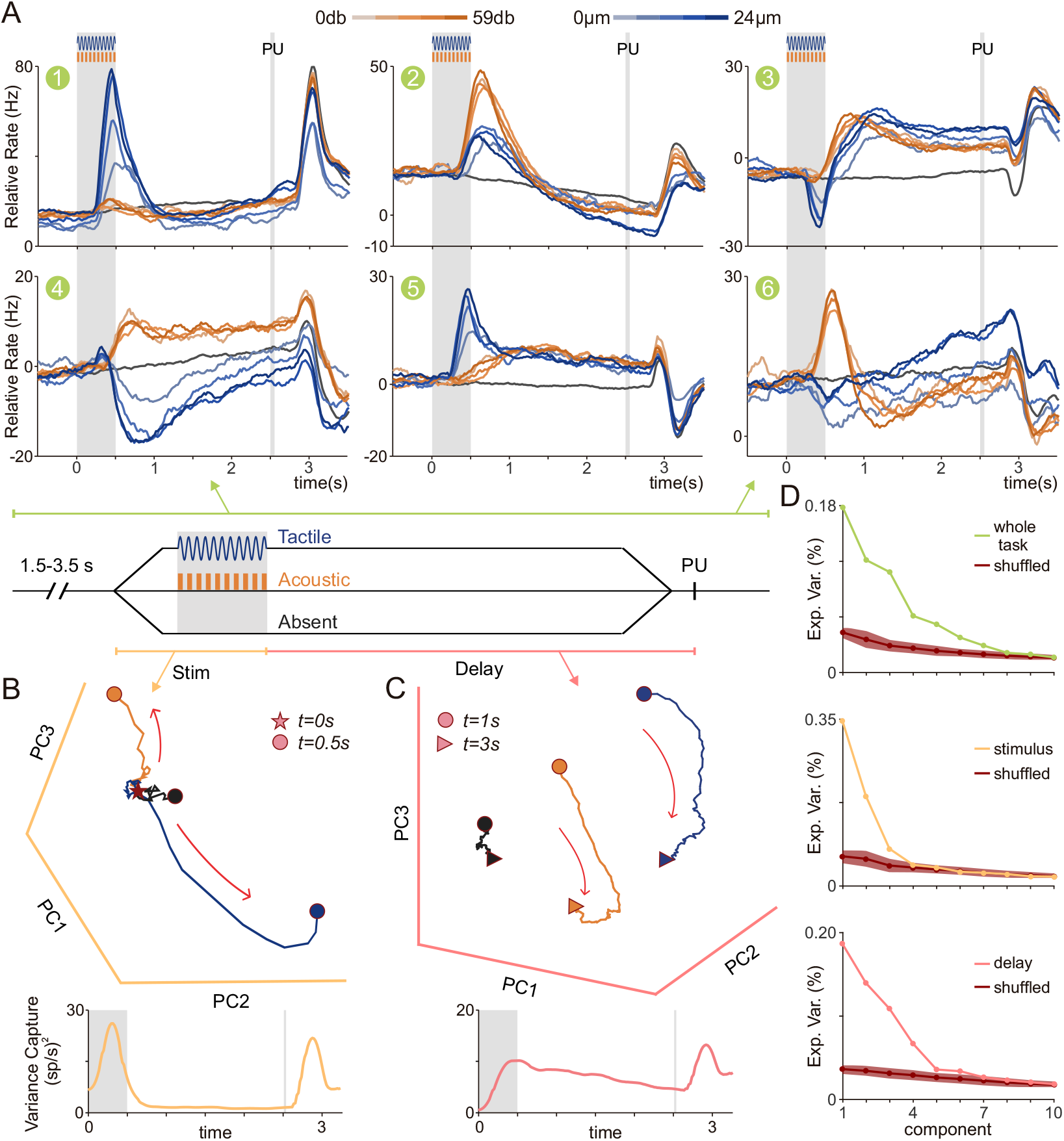
(A) First six PCs (EV, PC1: 17.8%; PC2: 12.1%; PC3: 10.8%; PC4: 6.1%; PC5: 5.2%; PC6: 3.7%) obtained from PCA of the covariance matrices derived from neural activity throughout the BDT (−1 to 3.5 s, green line on top of BDT schematic). Neural activity sorted by sensory modality and class was projected onto these axes. Activity associated with tactile (acoustic) stimulation is depicted by the traces in different tones of blue (orange), with darker (lighter) tones indicating stronger (softer) stimulation. Activity during stimulus-absent trials is depicted by the dark gray traces. Similarly to Figs. 4B and 4C, (B) Top: similar to (A) but applying PCA on the covariance matrix from neural activity during the stimulation period (0 to 0.5 s, yellow trace on BDT scheme). Neural responses associated with the same period and averaged by decision (tactile, acoustic, and stimulus-absent, shown as blue, orange, and black traces, respectively), were projected onto the first three PCs (stars represent t = 0 s while filled dots t = 0.5 s). Bottom: variance over time captured by the three PCs derived from the stimulation period. (C) Top: similar to (B) but PCs were derived from the covariance matrix using neural data from the delay period (0.5 to 1.5 s, pink line in the BDT schematic). Neural activity between 1 to 2.5 s of the BDT, averaged by decision, was then projected onto the first three PCs (filled dots represent t = 1 s while filled triangles t = 2.5 s). Bottom: variance over time captured by the three PCs derived in the delay period. (D) Explained variance of the first 10 PCs derived in (A) top, (B) middle, and (C) bottom. The dark-red traces, along with the standard error in each plot, represent the explained variance of the principal components (PCs) derived from applying PCA to shuffled covariance matrices, which were used to assess significance.

Restricting the analysis to the sensory period (Fig. 4B) yielded three significant PCs (Fig. 4D, middle). Projections onto these PCs revealed clear divergence of neural trajectories according to presented modality (tactile, acoustic, or absent), with tactile trials diverging more rapidly (Fig. S4F). Similar dynamics were observed when using full-task PCs for the stimulation period (Fig. S3A), consistent with the higher variance noted during sensory processing (Fig. 3A).

Focusing on the delay period (Fig. 4C), projection onto the first three significant delay-related PCs (Fig. 4D, bottom) revealed that trajectories, although initially segregated, remained distinct throughout the delay. This finding indicates that while sensory PCs capture dynamics during stimulation, delay-related axes reflect persistent, decision-related signals. Further analysis of mnemonic signals using established working memory methods (*33, 40*) (Fig. S3B) confirmed that these population signals remained stably segregated during the delay.

### Orthogonality between sensory and decision mnemonic dynamics

To further examine VPC population dynamics during stimulation and delay, we applied a generalized linear model (GLM) to the acoustic and tactile responses (Fig. 5). As before, tactile and acoustic trajectories diverged during stimulation, aligning with their respective coding axes (Fig. 5A). Similar results were obtained using sensory coefficients from a multi-linear model (MLM) (*41–43*). Notably, the sensory axes encoding amplitude from each modality evolved in nearly orthogonal directions (Fig. S3C), a finding confirmed by a simplified approach (*43, 44*) based on the difference between supra-threshold and absent activity (Fig. S3D). These results indicate that a simple linear decoder can reliably predict the presence of either acoustic or tactile stimulation from VPC neural activity.

**Figure 5.**
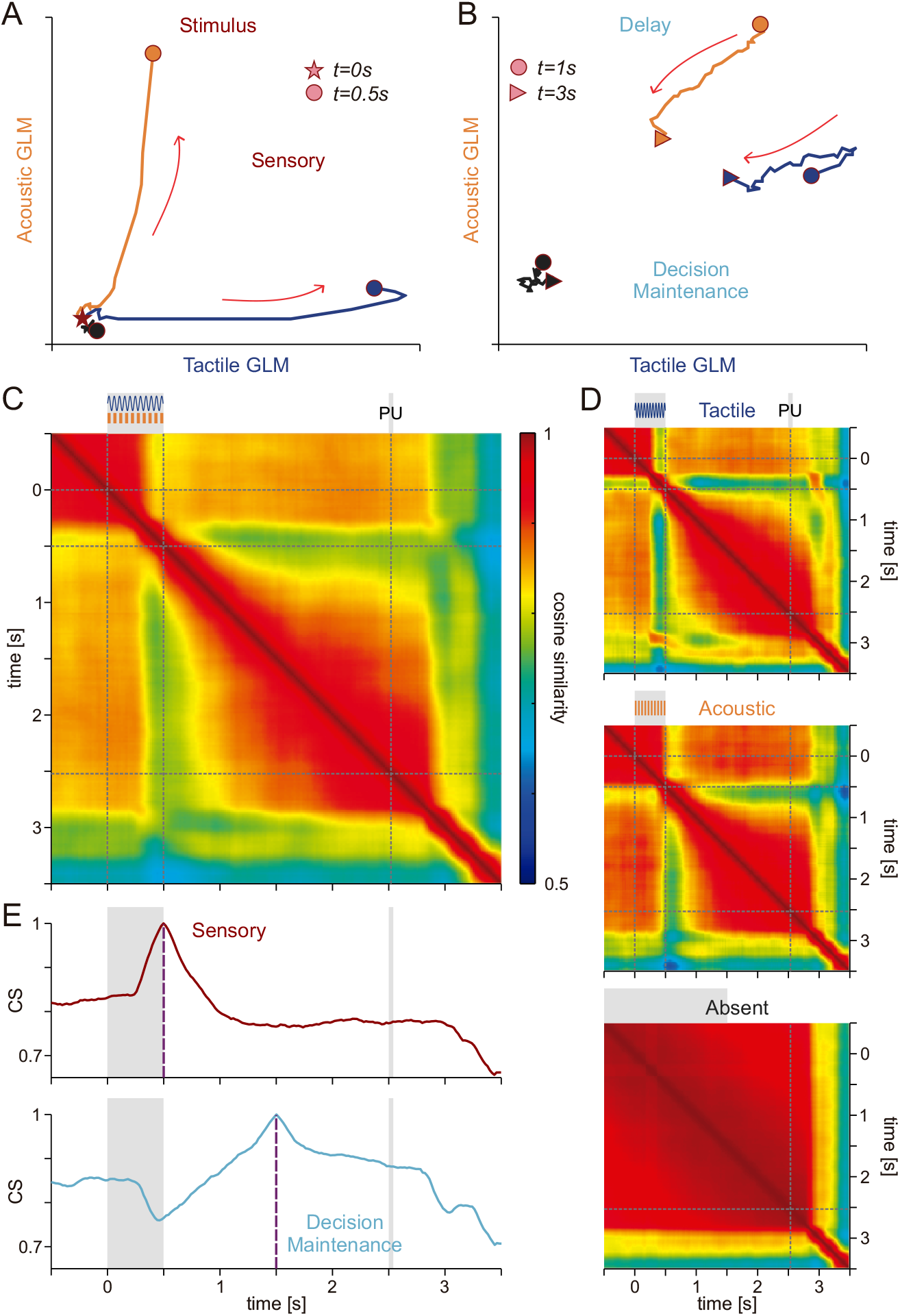
(A, B) Neural activity sorted and averaged by sensory modality was projected over the axes of a Generalized Linear Model (GLM) computed in the stimulus period (0 to 0.5 s, filled stars and dots mark start and end, respectively) (A), and in the delay period (1 to 2.5 s, filled stars, and triangles denote start and end, respectively) (B). Similarly to Figs 4B and 4C, activity for tactile, acoustic, and stimulus-absent decision are shown as blue, orange, and black traces, respectively (C, D) Cosine similarity matrix of the population states obtained by grouping (C) neural data from different modalities or sorted (D) by sensory stimulation (tactile on top, acoustic in the middle) and stimulus-absence (bottom). Dotted lines in each matrix indicate the stimulation period and PU event. Red and blue tones indicate high and low similarity among populational states, respectively. (E) Sensory (Top) and decision-maintenance (Bottom) cosine similarity over time computed using population vectors at t = 0.5 s (stimulation epoch) and 1.5 s (delay epoch), respectively.

During the delay period, neural trajectories remained distinct, though they converged slightly–likely due to reduced variance (Fig. 5B). Absent trials, in contrast, maintained a stable state throughout both the stimulus and delay periods, consistent with earlier observations (Figs. 4C and S3B).

Next we analyzed the similarity of population states throughout the task by calculating the cosine similarity (CS) between population activity vectors at different time points for each stimulus condition (*40, 45*). We averaged these similarities across all conditions (Fig. 5C), as well as separately for tactile (Fig. 5D top), acoustic (Fig. 5D middle), and absent trials (Fig. 5D bottom). This analysis revealed a clear dissociation between sensory processing and the remainder of the task: the VPC population exhibited a distinct state during sensory arrival, which then stabilized decision delay. Sorting trials by modality confirmed that tactile responses emerged faster than acoustic responses (in line with Fig. 3G), and that the tactile sensory closely resemble the network state after the “pu” signal.

Absent trials, despite some neurons showing ramping activity (Figs. 1C, D; S1A, G, E, I), consistently maintained a high similarity throughout the task. This suggests that while ramping modulates firing rate magnitude, it does not alter the overall direction–or “angle”–of the population vector, preserving a stable representation in the absence of sensory input.

Finally, Fig. 5E presents the CS across all time points, highlighting two key moments: during the sensory period (t = 0.5s) and during the memory maintenance delay (t = 1.5s). Consistent with the sensory versus memory decoupling observed in Fig. 4B (bottom), the population state during sensory processing is distinct from that during delay, as evidenced by a green band (t = 0.3 and t = 0.7 s) in the *TxT* matrix (Fig. 5C) and a separate, persistent red rhomboid-shaped pattern during the delay.

### Rotational Dynamics from Stimulation to Decision Maintenance

To disentangle condition-averaged temporal dynamics from modality-specific responses, we segregated neural activity into modality-dependent and modality-independent components. Prior studies have shown that condition independent signals account for much of the network’s response (*32, 39, 46*). To achieve this, we marginalized the covariance matrices into modal and amodal parts and then we applied demixed principal component analysis (dPCA) (*32*) to extract the latent responses corresponding to each. In Fig. 6A, the first six dPCA components are shown, ordered by explained variance (Fig. 6B). The first three components–capturing most of the variance–were amodal, reflecting dynamics during both stimulation and response periods regardless of stimulus modality, and they also exhibited ramping during absent trials, consistent with earlier observations.

**Figure 6.**
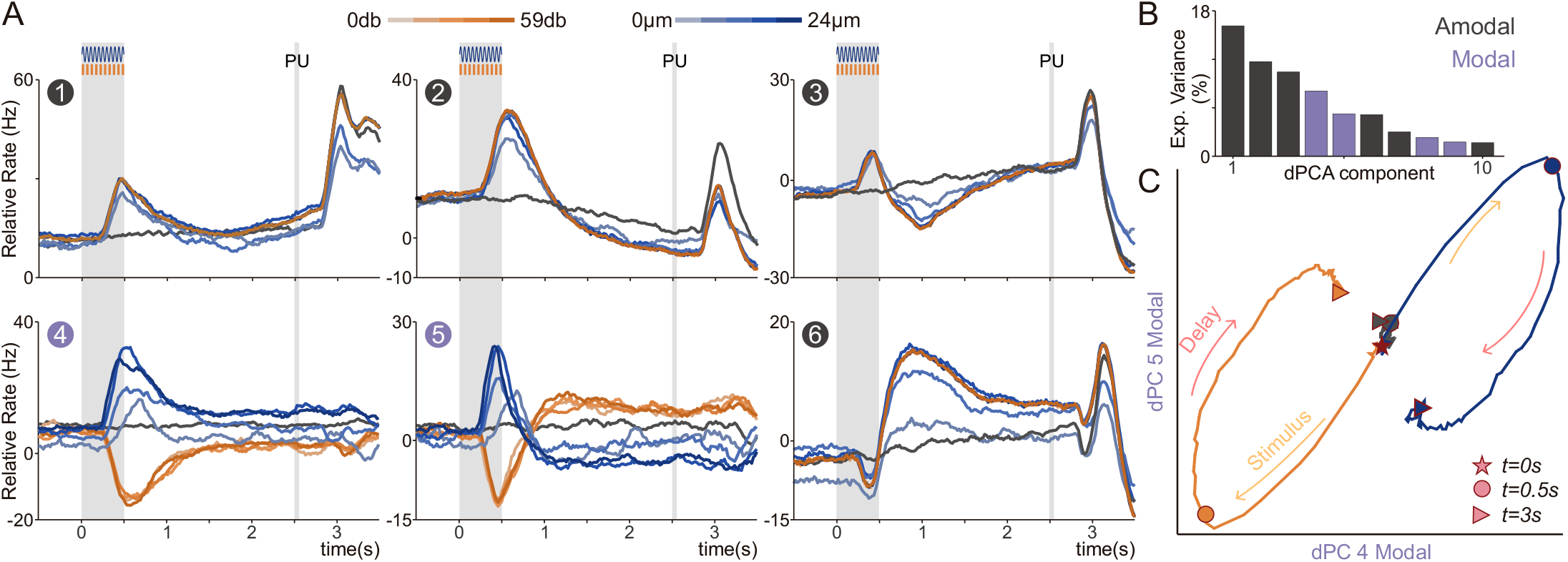
(A) Demixed principal component analysis (dPCA) applied to marginalized covariance matrices from the whole BDT (−1 to 3.5 s) with respect to class identity (tactile, acoustic) to derive the most prominent dPCs. Population activity for the entire BDT (−1 to 3.5 s, n=911), sorted by class averages within each sensory modality (tactile and acoustic, tones of blue and orange traces, respectively) or absence of stimulation (absent, dark-gray trace) projected onto the first six dPCs ordered by their EV (dPC1: 16.1%, dPC2: 11.6%, dPC3: 10.3%, dPC4: 8.05%, dPC5: 5.2%, dPC6: 5.1%). Projection onto most dPCs showed amodal coding behavior, while projections onto dPC4 and dPC5 displayed modal dynamics with clear tactile, acoustic, and stimulus-absent encoding and maintenance. Gray-shaded rectangles represent the stimulation period and PU. (B) EV of the first 10 dPCs with modal and amodal dPCs represented by purple and gray rectangles, respectively. (C) Neural activity across the whole task (0 to 3.5 s) sorted by modality (tactile [blue], acoustic [orange], and stimulus-absent [black]) projected over the first two modal dPCs (dPC4 & dPC5) and plotted in a 2D phase diagram. Filled stars, dots, and triangles mark moments t=0 s, 0.5 s, and 2.5 s, respectively.

In contrast, the next two components were modality-specific, capturing the bulk of the dynamics during stimulus arrival and decision maintenance. These modal components were active during stimulation and the decision delay but not after the “pu” period. When projecting the population activity onto these two components (Fig. 6C), we observed a clear separation of tactile and acoustic trajectories during stimulation, followed by a rapid rotation at the beginning of the delay and a slow convergence toward a subspace orthogonal to the initial sensory response. These findings suggest that the VPC population exhibits rotational dynamics that may help reduce interference between sensory inputs and decision representations–a mechanism previously observed in rodents during learning to minimize conflicts between sensory and mnemonic signals (*28*).

### Common Multistability Mechanism Across Artificial and Biological Networks

We investigated whether the dynamics observed in the VPC could be compared to those in a recurrent neural network (RNN) trained to solve the BDT (Fig. 7A). This network architecture has been successfully employed to reproduce population dynamics during other cognitive tasks (*30, 31, 41, 47, 48*) and can switch among many previously learned cognitive tasks (*48*). Notably, these models exhibited comparable dynamics to biological networks (*41, 43, 47, 49, 50*). We recreated our BDT in the model with two input nodes, each representing a sensory channel of a distinct modality, and three output nodes, corresponding to the response buttons in the macaques’ task. Following standard methodology (*30*), the network was trained until it reached good performance (Methods, Fig.7B, C).

**Figure 7.**
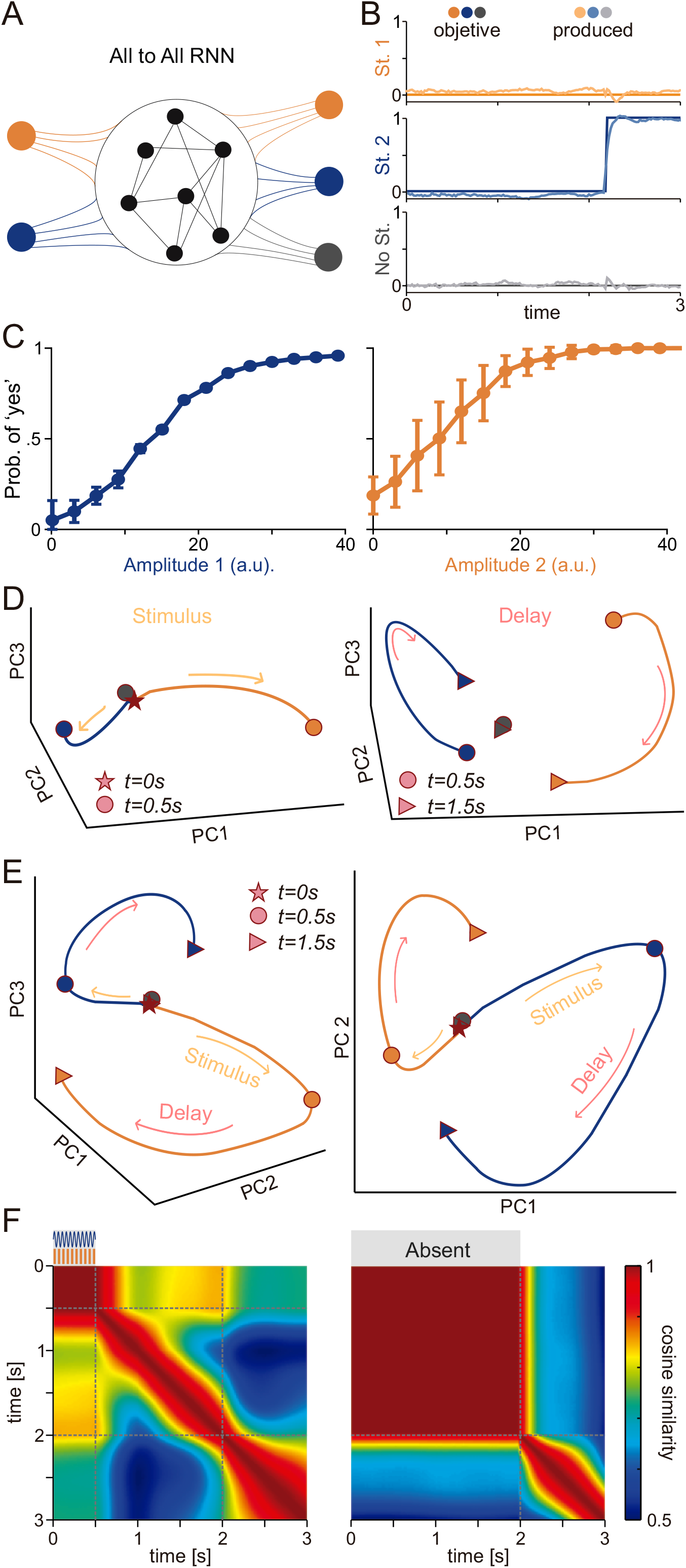
(A) Schematic representation of an all-to-all RNN trained for BDT. The network processes input stimuli—stimulus 1 (St. 1, tactile, blue) and stimulus 2 (St. 2, acoustic, orange) are implemented as the two nodes on the left—to generate corresponding decisional outputs: tactile, acoustic, or absent stimuli, implemented as the blue, orange, and gray nodes on the right, respectively. (B) Examples of three supra-liminal trials after RNN training for St.1 (tactile, blue traces), St. 2 (acoustic, orange traces), and No St. (absent, grey traces). Traces in saturated and unsaturated tones correspond to the objective and produced outputs, respectively. After sensory input presentation (St.1, St. 2, or No St.), the RNN is trained to maintain quiescence until the PU event (go-cue), when the activity of the output channel associated with the presented stimulus class is expected to rise (orange and blue for St.1 and St.2, respectively; gray for No St. - shown here for St. 2). (C) Post-training psychometric curves corresponding to each stimulus type. The training utilized a limited range of possible stimulus amplitudes. (D) PCA applied to covariance matrices obtained using RNN neural activity from the stimulus (0 to 0.5 s) and decision (0.5 to 1.5 s) epochs to derive the most prominent PCs. Simulated neural data during the stimulus (0 to 0.5 s, panel D, left) or delay (0.5 to 1.5 s, panel D, right) periods, sorted by modality (tactile [blue], acoustic [orange], and absent [black]), projected onto the first three PCs. RNN network dynamics displayed outstanding similarity with biological data. (E) PCA applied to the activity from the whole simulated BDT without the response period (0 to 2 s). Dynamics diverge according to sensory input (3D diagram on the left), similarly to those obtained by the modal axes dPC4 & 5 (2D diagram on the right). (F) Cosine similarity matrix of the populational states computed using trials from all modalities (tactile, acoustic, and stimulus-absent, matrix on the left) or solely using stimulus-absent trials (matrix on the right). Computed cosine similarity matrices are reminiscent of those computed using biological neural activity. The dotted lines indicate the stimulus period and the PU event.

In concordance with VPC population responses (Fig. 4B), the RNN showed a divergence of population trajectories during stimulation (Fig. 7D left). Sensory representation was followed by separated and rotational tactile and acoustic responses during the delay period (Fig. 7D right and 7E), similar to what was observed in the VPC (Fig. 6C). Similarly to the biological network, in absent trials the population response did not change its angle until the response onset (Fig. 7F). The similar dynamics found in the RNN model and the VPC networks suggest that the mechanisms studied in the VPC allow for general bimodal information integration and bimodal perceptual decision maintenance.

### A general dynamical mechanism for the transition from sensory to mnemonic representations

To propose a general mechanism underlying the results obtained with biological and artificial networks, we derived a low-dimensional dynamical system model that captures the separation between sensory and maintenance axes (Fig. 8A). The proposed dynamical system (Eq. S17 and S18 and Fig. 8B) includes two saddle points whose stable manifolds act as sensory thresholds for detecting each sensory modality. When the stimulus intensity exceeds a certain threshold, the acoustic (orange) and tactile (blue) responses cross the respective sensory threshold (green line) and subsequently relax toward stable equilibrium points associated with memory retention for the two sensory modalities. However, if the stimulus is too weak, the response fails to reach the threshold and returns to the stable equilibrium associated with the absence of sensory input (Fig. 8C). See also Fig. S5 for the model’s response to increasingly stronger tactile and acoustic stimuli.

**Figure 8.**
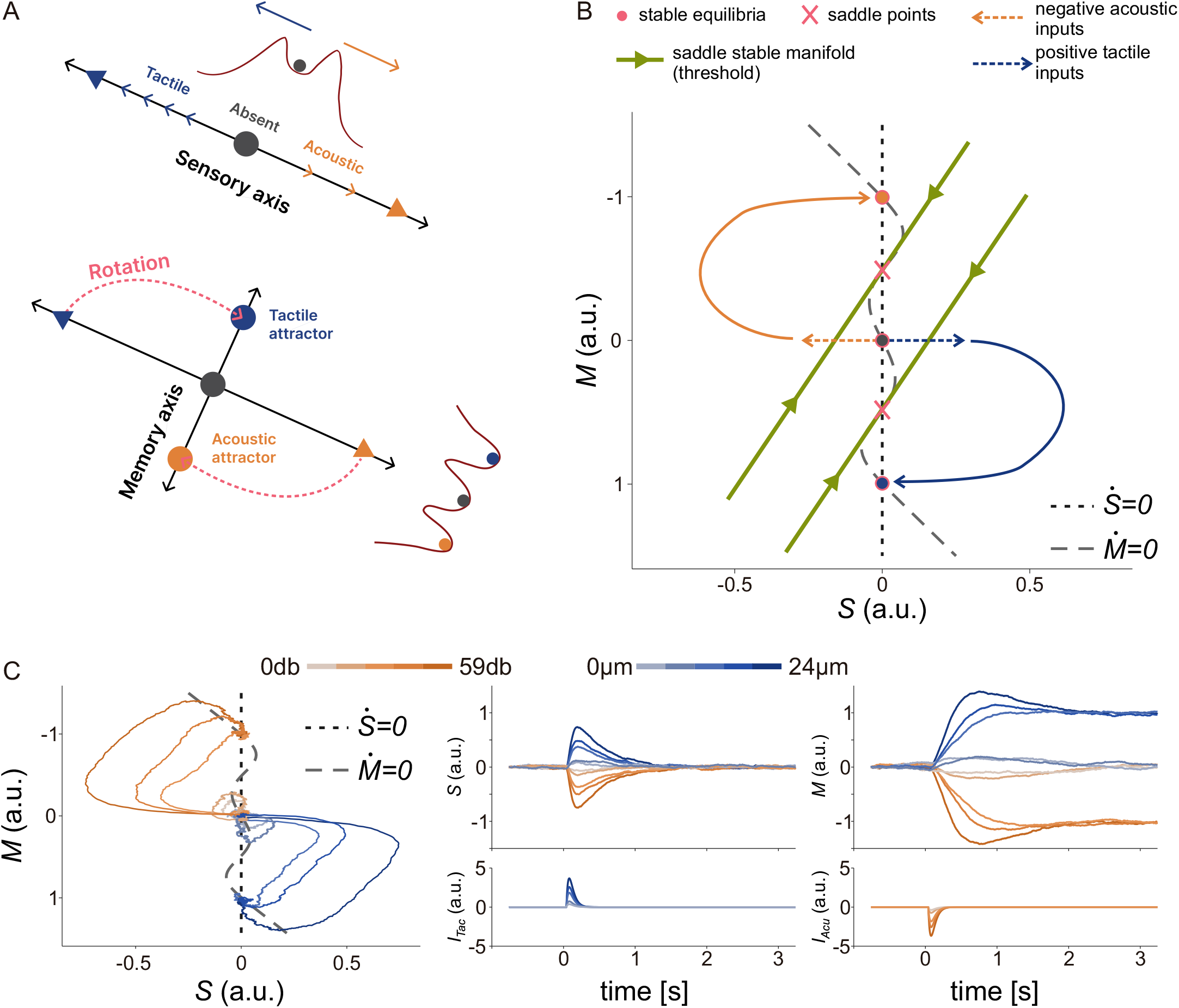
(A) Sketch of a low-dimensional dynamics performing bimodal detection. When receiving a sufficiently strong stimulus from one of the two modalities modality, the dynamics integrates it along the sensory axis. After the stimulation ends, the state converges with a rotational motion toward a memory subspace that preserves the memory of the stimulus’s presence and modality in distinct attractors. (B) Sketch of the phase plane of the suggested bimodal detection dynamics. The fixed points of the system are represented as filled pink circles for stable equilibria and crosses for saddle points. The S and M nullclines are depicted with dotted and dashed gray lines, respectively. Saddle stable manifolds are sketched in green. (C) Example runs for three supra-threshold stimuli and two sub-threshold stimuli of distinct amplitudes. Left: System’s phase plane. Middle: Sensory integration variable over time. Right: Memory retention variable over time. Bottom: Sensory input used for the tactile and acoustic runs. The proposed two-dimensional model qualitatively reproduces and explains the dynamics shown in Figures 6C and 7E.

The reduced model in Eqs. S17 and S18 provides a system-level description of the dynamical interaction between sensory and mnemonic representations. According to this model, sensory representation dynamics (Eq. S17) act as a simple linear low-pass filter on sensory inputs. The output of these sensory dynamics feeds into the mnemonic representation dynamics (Eq. S18), which are nonlinear and exhibit multi-stability among attractors corresponding to distinct perceptual decisions. The rotational geometry observed during the transition from sensory to mnemonic representation (Fig. 6C and Fig. 7E) is explained by the phase-plane geometry of the sensory-mnemonic dynamical system described by equations by Eqs. S17 and S18: a cascade of a vanishing-memory low-pass filter for transient sensory input representation, followed by a nonlinear multi-stable system for persistent memory retention of relevant sensory input features.

Due to the universality of low-pass filtering and multi-stable dynamics, the proposed model offers a general dynamical mechanism likely shared across sensory modalities and animal model tasks. This mechanism elucidates the transition from sensory to mnemonic representations within neural populations, contributing to a deeper understanding of how sensory information is retained and transformed into memory across different contexts.

## DISCUSSION

Our investigation of the BDT revealed that the VPC serves as a natural integrator of tactile and acoustic sensory inputs. Neurons in this region exhibited diverse response patterns, encoding stimulus presence, absence and modality. Tactile responses emerged first, but the VPC consistently maintained information across sensory modalities, underscoring its role in integrating multimodal inputs for decision-making. These findings, replicated across subjects and hemispheres, highlight the VPC’s contribution to sensory differentiation and cognitive processes.

Population-level analyses distinguished sensory responses from decision maintenance, revealing low-dimensional neural trajectories that initially diverged upon stimulation before rotating into a subspace orthogonal to the sensory response. This same behavior was replicated in an RNN trained on the BDT, supporting the hypothesis of a shared computational mechanism. Using a simplify two-dimensional model, we demonstrated that when sensory input was sufficiently strong, the system crossed a separatrix, stabilizing in an attractor state that maintained the decision. This suggests the VPC is not only involved in sensory encoding but also in sustaining decisions through distinct neural mechanisms.

The separation between sensory and maintenance dynamics is crucial because it allows the brain to retain information using a different coding scheme than that used during sensory reception. This strategy enables continuous processing and attention to new stimuli without disrupting the maintenance of previously encoded information (*28*) or impairing communication with other brain areas (*51*). Our work thus reveals a novel bimodal mechanism in the VPC that provides unique insight into how two competing sensory modalities can be integrated.

Furthermore, it is important to consider how sensory integration in the VPC might change under constructive interference, rather than competition. Constructive interference could enhance the fidelity and robustness of encoded information, potentially leading to more accurate and faster decision-making processes (*12, 52–54*). Such interference might allow the VPC to combine inputs from different sensory modalities in a manner that strengthens the overall signal, facilitating clearer perceptual decisions. Understanding whether and how VPC leverages constructive interference will be essential for clarifying its role in multimodal sensory processing.

Several questions remain. For example, why does tactile encoding emerge faster than acoustic encoding? One possibility is that the task’s heavy reliance on tactile input triggers a rapid response to touch, or that non-human primates are less adept at processing acoustic stimuli. Unlike primary sensory areas (S1 (*17*) and A1 (*13*)), the VPC exhibits bimodal encoding, raising the possibility that it might be the first cortical area to feature neurons with distinct modality-specific responses (*2, 19*). While not conclusive, the VPC’s rapid and robust responses make it a strong candidate for this role.

Recent research has also implicated the VPC in abstract language processing. Traditionally, Broca’s area has been considered the hub for language, but emerging evidence suggests that the VPC contributes significantly to language-related representations (*55*). Language is inherently multimodal–integrating acoustic, visual, and tactile information (*56*)–and our findings support the notion that the VPC is vital role for encoding and maintaining such multimodal sensory information, a capacity that could be critical for higher-order cognitive functions like language.

The role of oscillatory activity in bimodal integration remains an open question. It is possible that the bimodal coding we observed is reflected in local field potential frequency bands, with the beta band being an strong candidate (*57, 58*). Although the specific role of beta oscillations in the VPC is not fully understood, their established role in other brain regions suggests they could facilitate communication between different sensory areas and the VPC through spikes synchronization (*59*). Such synchronization may regulate sensory attention and enhance the integration of task-relevant inputs.

Attention also appears to modulate VPC activity. A reasonable hypothesis is that neural responses in the VPC are attention-dependent: if a sensory modality becomes irrelevant, the corresponding neural activity may decrease or cease (*41, 60*). Future experiments that vary attention could elucidate how VPC neurons adjust their responses based on context, deepening our understanding of sensory integration and attention regulation.

In summary, when compared with previous studies (*2, 19, 26, 27, 60*), our findings demonstrate that the VPC plays a key role in segregating sensory inputs and sustaining multimodal perceptual alternatives. The pronounced segregation of sensory and maintenance responses within a low-dimensional neural space–in both biological and artificial systems–illustrates the versatility of VPC dynamics. This research not only shows that a mechanism previously thought to be restricted to unimodal contexts is effective in bimodal conditions but also paves the way for further investigation into how the brain processes and retains multisensory information.

## ACKNOWLEDGMENTS

This work was supported by grants PAPIIT-IN210819 (to R.R. and R.R.-P.) and PAPIIT-IN205022 (to R.R.-P.) and IN203825 (to R.R.-P.) from the Dirección de Asuntos del Personal Académico de la Universidad Nacional Autónoma de México and CONAHCYT-240892 (to R.R.), CONAHCYT-319347 (to R.R.-P.) from Consejo Nacional de Ciencia y Tecnología; IBRO Early Career Award 2022 (to R.R.-P.) from International Brain Research Association. H.D. is a doctoral student from Programa de Doctorado en Ciencias Biomédicas, UNAM. L.B. is a postdoctoral researcher (Postdoctoral fellowship CONACYT-838783).

## Methods

### Bimodal Detection Task

Two monkeys (*Macaca mulatta*) were trained to detect the presence or absence of either a vibrotactile stimulus or acoustic stimulus (Fig. 1A) (*17*). Vibrotactile stimuli ranged from 0 to 24 µm, while acoustic stimuli ranged from 0 to 59 dB, covering sub-threshold, threshold, and supra-threshold detection levels for both modalities. The vibrotactile stimuli were delivered via a computer-controlled stimulator with a 2-mm round tip (BME Systems) applied to the glabrous skin of the distal segment of one fingertip on the monkey’s restrained right hand, while the acoustic stimuli were presented through a speaker. The task began when the stimulator probe (PD) was lowered onto the finger’s surface, signaling the start of a trial (PD). The monkey responded by grasping the lever (KD). After a variable delay (indicate in Fig. 1A), the monkey received either a tactile stimulus in one-third of the trials, an acoustic stimulus in another third, or no stimulus at all in the remaining third. After a 2-second delay, the stimulator was lifted off from the fingertip (PU), signaling the monkey to release the lever (KU) and press a button (PB) to indicate the perceived stimulus modality or its absence. Both types of stimuli consisted of sinusoidal signals at a frequency of 20 Hz. For the acoustic modality, each signal was a pure tone at 1 kHz, while for the vibrotactile modality, the stimuli were trains of mechanical-sinusoidal pulses. The trials alternated between stimuli and no-stimulus conditions in equal proportions. The monkeys indicated their detection by pressing one of three buttons: “vibrotactile stimulus present,” “acoustic stimulus present,” or “stimulus absent.” Correct responses were rewarded with a few drops of fruit juice. Performance was quantified using psychometric techniques (Fig. 1E). Both animals were handled in accordance with the institutional standards of the U.S. National Institutes of Health and the Society for Neuroscience. All protocols were approved by the Institutional Animal Care and Use Committee of the Instituto de Fisiología Celular at the National Autonomous University of Mexico (UNAM).

### Recording sites and data sets

Neuronal recordings were obtained with an array of seven independent, movable microelectrodes (2–3 [MΩ]) inserted into VPC (Fig. 1B) (*20–22, 26*). In monkey RR32, recordings were made contralaterally (right hemisphere) and ipsilaterally (left hemisphere) to the stimulated hand, while in monkey RR33, recordings were made only ipsilaterally to motor execution (right hemisphere). The electrode locations were confirmed through standard histological techniques. Each session included an equal number of trials for tactile (50), acoustic (50), and absent (50) stimuli. Recording sites varied across sessions, with each set of sites lasting for approximately 30 sessions (1 session per day). A total of 42 recording sessions were conducted for monkey RR32, and 27 for RR33. The electrode penetration locations were mapped onto the cortical surface by marking the edges of the small chamber (7 mm in diameter) placed over each cortical area. Across the sessions, we recorded 566 neurons in monkey RR32 (256 from the right hemisphere and 310 from the left hemisphere) and 345 neurons in RR33. It is important to note that neurons displaying a preference for encoding one of the modalities (tactile or acoustic) were distributed throughout the VPC, making the identification of subgroup responses highly unlikely (see Fig. 3E and Fig. S2, A–C, top and middle panels).

### Data analysis

For the analyses described below and the results presented in this study, only hit trials were considered, and subliminal classes were excluded when plotting the class averages. This approach ensures that the data reflects accurate sensory detection and response, focusing solely on trials where the monkeys successfully identified the presence of stimuli.

#### Firing Rate

For each neuron, we calculated a time-dependent firing rate for each trial using overlapping rectangular causal windows with a length of 200 ms and steps of 20 ms. For visualization purposes only, as shown in Figs. 2 and S1, we applied a causal Gaussian convolution with a σ of 200 ms. This smoothing technique allows for a clearer representation of firing rate dynamics over time, without distorting the temporal relationships in the neuronal activity.

#### Instantaneous coding variance across the population

To evaluate the influence of each type of sensory stimulus coding (tactile or acoustic) and decision outcome, we calculated the instantaneous variance in a time-dependent manner using the computed firing rate of each neuron. At each time bin, the population’s instantaneous tactile variance (*Var*_*Tac*_, Figs. 3A & S2 middle panels from A to C, blue trace in all plots) was calculated as the quadratic sum of the firing rate fluctuations across different tactile stimulus identities (classes) and neurons:

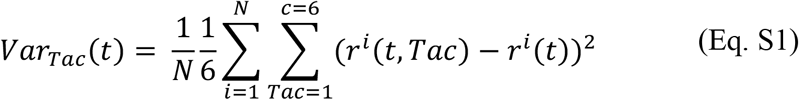

This metric captures the variability in neural responses specifically linked to the tactile stimuli at any given moment in time, providing insight into how the population dynamically encodes this sensory information.

Analogously, the population’s instantaneous acoustic variance (*Var*_*Acu*_, Figs. 3A & S2 middle panels from A to C, orange trace in all plots) was calculated to measure the fluctuations in firing rate across acoustic stimulus classes and neurons:

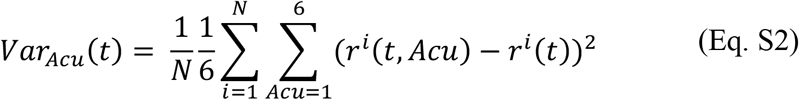

Next, the population’s instantaneous decision variance (*Var*_*Dec*_, Figs. 3A & S2 middle panels from A to C, purple trace in all plots) was calculated to measure the fluctuations in firing rate across different decision identities and neurons:

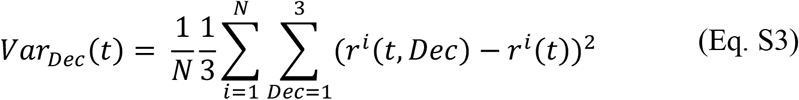

This metric quantifies the variability in neural responses associated with the different decision outcomes over time.

The tactile and acoustic variances were obtained by using only the means of the stimulus classes within their respective modalities. The decision variance, on the other hand, was calculated by averaging across all classes for each decision type (Tactile, Acoustic, No Stimulus). The overall class variance, which reflects variability across the entire range of stimuli, was computed using all classes, including all amplitudes for acoustic, tactile, and no stimulus trials.

The values of *Var*_*Tac*_, *Var*_*Acu*_, and *Var*_*Dec*_, computed from the activity within the time window t = -1 to 0s (before the arrival (or not) of stimulation), represent the inherent residual noise in the firing rate estimates (∼2 spikes/second). To interpret these values as a measure of population coding, *Var*_*Tac*_, *Var*_*Acu*_, and *Var*_*Dec*_ must exceed this resting state variance (basal variance). In our analysis, to ensure meaningful comparisons across parameters, the basal variance was subtracted from each type of variance computed. This adjustment allows us to more accurately assess the degree of coding attributable to sensory stimuli and decision processes, beyond the inherent fluctuations.

#### Instantaneous temporal variance across the population

The population instantaneous temporal variance (*Var*_*Temp*_, Figs. 3B & S2 bottom panels from A to C, green trace in all plots) was computed at each time point as compared to the mean firing rate. This variance is a reflection of the time variation with respect to the baseline period. In particular, it is defined as the quadratic sum of differences between the average firing rate at each time point (*r*^*i*^ (*t*)) and the average baseline firing rate 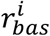 (from -1 to 0 s) across neurons:

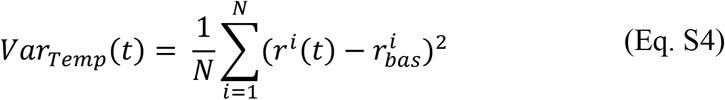

This metric gives an account of the dynamics in the population firing rate over time relative to its baseline activity, emphasizing the temporal dynamics in the neural population throughout the task.

#### Mutual Information

For each neuron, we calculated four types of mutual information for each time window using Shannon’s information theory (*61, 62*): information modality coding (I_COD_, depicted as Cod in Fig. 3C) and information derived from the combinations of possible decisions: Tactile-Acoustic (TA), Tactile-No Stimulus (NT), and Acoustic-No Stimulus (NA). To compute *I*_*COD*_, we considered the firing rate probability distributions associated with the different stimulus classes within each modality (*P*(*r*|*s*)) as well as the overall firing rate probability distribution (*P*(*r*)), which combines all stimulus classes and modalities. For each time bin, we generated three conditional distributions *P*(*r*|*s*): one accounted for each one of the tactile classes (*P*(*r*|*Tac*)), one for each of the acoustic classes (*P*(*r*|*Acu*)), and one for stimulus absent class (*P*(*r*|*Abs*). Additionally, we computed the unconditioned probability distribution (*P*(*r*)), which reflects the general likelihood of any neural response across all classes and modalities. The coding information was then quantified using mutual information:

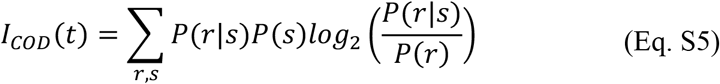

This measures the amount of information the neural response *r* provides about the stimulus *s*.

To compute the information metric for the combinations *I*_*TA*_, *I*_*NT*_, and *I*_*NA*_, we followed the same procedure as for *I*_*COD*_ (Eq. S5) but utilized the corresponding firing rate probability distributions *P*(*r*|*s*) based on the trials of stimuli classes and modalities involved in each pair of combinations. For each time bin, we generated three distributions (same as *I*_*COD*_): tactile and acoustic distributions for *I*_*TA*_; tactile and absent distributions for *I*_*NT*_; and acoustic and absent distributions for *I*_*NA*_ (*17*). In all cases, the unconditioned probability distribution *P*(*r*) was computed (in each time) by combining the relevant trials from the stimuli classes and modalities. Neurons were then categorized (as shown in the chart in Fig. 3D) based on whether they showed significant information for at least two out of the three possible information metrics (*I*_*TA*_, *I*_*NT*_, *I*_*NA*_). They were classified as: Decisional (Dec) if they scored significantly in all combinations (*I*_*TA*_, *I*_*NT*_, *I*_*NA*_); Tactile (Tac) if they scored in *I*_*TA*_ and *I*_*NT*_; Acoustic (Acu) if they scored in *I*_*TA*_ and *I*_*NA*_; and finally Absent (No) if they carried relevant information for the *I*_*NT*_ and *I*_*NA*_ pair. The sorted neuron subgroups were plotted as the percentage of neurons over time (represented by traces and pie charts in Fig. 3E). The pie chart considered the total number of significant neurons. Neurons that showed significant information for only one of the possible combinations were not classified into a specific group but were included in the total neuron count. This subgroup is shown as the white portions of the pie charts (Fig. 3E).

For all information metrics computed, significance was calculated per neuron in each time bin. A permuted value of mutual information was obtained by performing 1,000 permutations of the trials. A mutual information value was considered significant if the probability of a permutation yielding the same or a greater value was below 0.05 (p < 0.05). The percentage of neurons with significant mutual information was then corrected using a multiple comparison test to account for the number of neurons and time bins analyzed. Additionally, to address the issue of finite sampling, we applied the correction proposed in^54^. For the significance test, we also applied a correction for multiple comparisons using a clustering method as described in (*63*), retaining only the sets of significance-connected time bins whose size exceeded a defined threshold.

#### Coding latency

This metric denotes the time at which the encoded signals first become significant (*2*). We calculated coding latencies for the four subgroups of categorized neurons (Dec, Tac, Acu, and No). For each subgroup, latency was defined as the first time window (after stimulus presentation) that was followed by three consecutive time windows with a significant p-value. To determine the number of consecutive bins (3 in this case) required to establish significance, we applied a significance-by-size approach, ensuring that the cluster size was unlikely to occur by chance (p < 0.05).

#### Population States Similarity

To analyze the similarity of population states across different times of the task (*t*_1_, *t*_2_), within each modality (tactile or acoustic, Fig. 5D top & middle panels) or for the absence of stimulation (stimulus absent, Fig. 5D bottom panel), we computed a population vector for each time window of the BDT. The vector takes the form:

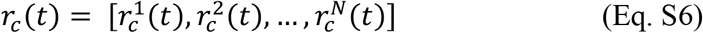

where each element of this vector corresponds to the mean firing rate of all trials for each neuron *i* under condition *c*, at the specific time 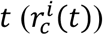. This vector, *r*_*c*_ (*t*), captures the population activity patterns across neurons for a given condition at different time points. Additionally, to compare the population states across conditions (modalities and amplitudes) (Fig. 5C), a total population vector was calculated by concatenating the vectors for each condition *c* at each time window *t*:

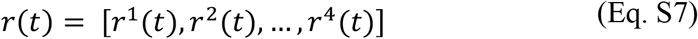

This represents the combined neural activity from all conditions and neurons, allowing for direct comparison of population states across different stimulus classes and modalities at various time points. The comparison between population states was assessed by computing the cosine similarity metric (*33, 40, 45*). This measure calculates the cosine of the angle between two vectors, without considering their magnitudes. The cosine similarity (*CS*) is 1 for two parallel vectors and 0 for orthogonal vectors (90° difference). Cosine similarity can be mathematically represented as:

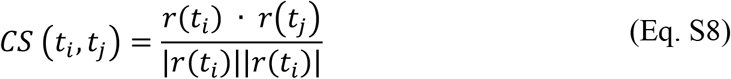

where *r*(*t*_*i*_) · *r*(*t*_j_) is the dot product of two vectors at times *t*_*i*_ and *t*_j_ respectively, and |*r*(*t*_*i*_)| and I*r*(*t*_j_)I are the magnitudes of the vectors at those same time points. Therefore, *CS* (*t*_*i*_, *t*_j_) defines a matrix of dimension *T*x *T*, which quantifies the similarity of population states at two different times. In this work, we quantified these matrices by considering all conditions and modalities (Figure 5C) or by separating the trials according to their modality (tactile, acoustic, or absent, Figure 5D). Analogous analyses were applied to the artificial networks in Figure 7.

#### Principal Component Analysis (PCA)

The main aim of PCA is to find a new coordinate system in which the data can be represented more succinctly and compactly. In other words, the idea is to define a low-dimensional subspace that captures most of the variance of the high-dimensional neural state space (*29, 39*). Typically, the number of significant dimensions is reduced substantially, going from as many possible dimensions as there are neurons (*N*) to just a few axes that concentrate most of the variance (*37*). To characterize how the population activity covaries across classes as a function of time, we performed PCA (Figs. 3 and S3 A & B) using liminal (4) and supraliminal (4) classes from the different stimulus modalities (tactile and acoustic) and the stimulus-absent condition (1 class), combining variance over classes and time (from -1 to 3.5 s for the whole BDT in Fig. 3A and S3A, and smaller periods in Figs. 3B & C and S3B). PCA yields a new coordinate system for the N-dimensional data, where the first coordinate accounts for the most variance in the neural population. The second coordinate accounts for as much of the remaining variance as possible, and so on, with each subsequent axis constrained to be orthogonal to the previous ones. PCA was computed from the firing rate covariance matrix, averaging over time bins *t* and classes *c*:

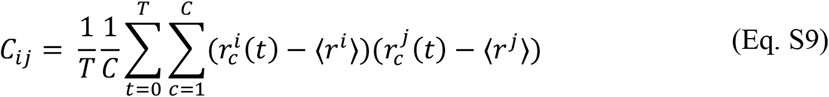

Where *C*_*ij*_ represents the covariance between neuron *i* and neuron *j, T* is the number of time windows in the period being considered, *C* is the number of conditions (9 in our task), 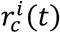 is the firing rate of neuron *i* at time *t* for class *c*, and ⟨*r*^*i*^⟩ is the mean firing rate of neuron *i* across all time bins and conditions. The diagonalization of the covariance matrix, *C* = *UDU*^*T*^, produces a new coordinate system given by the columns of matrix *U*, which we call the derived axes or principal components (PCs). On the other hand, *D* is a diagonal matrix of positive values. The diagonal elements of *D* represent the amount of variance in the population activity captured by the corresponding PCs. We then order the PCs according to the amount of variance captured. The projection of the N-dimensional data onto the *k*-th PC is given by:

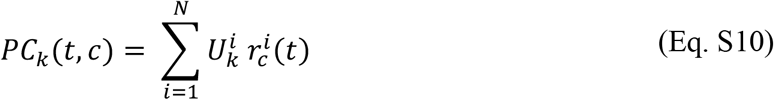

where 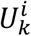 is the *i* element of the *k* PC (*U*_k_). Therefore, the PCs are linear readouts of the population activity; in other words, they are orthogonal linear combinations of the firing rates of the individual neurons, with no correlation between components (Fig. S4A-D). Thus, the contribution of each neuron to a given *PC*_k_ is given by the *ith* element of *U*_k_. These PCs can be thought of as a low-dimensional description of the population activity in this coding subspace. In Fig. 3 and S3 (A & B), principal components (PCs) were calculated for the BDT (n=911) by using neural activity from the entire task (−1 to 3.5 s of BDT, green line on task schematic in Fig. 3 and S3) or by using only data from the stimulation (0 to 0.5 s) or delay (0.5 to 2.5 s) periods (yellow and pink lines, respectively, on the task schematic in Fig. 3 and Fig. S3). Notably, the mnemonic PCs (mPCs) shown in Fig. S3B were computed by applying PCA to covariance matrices obtained from time-averaged neural population vectors during the delay period (0.5 to 2.5 s of BDT). For each experimental condition *c*, the elements for each neuron were averaged over the different time bins *i*, effectively collapsing the temporal dimension (*33, 40*)..

#### Speed of population dynamics

In order to calculate the speed of the population dynamics, we employed the embedding space of the first significant PCs. After projecting the trajectories of each class to this space, we measured their evolution at each time step with the euclidean distance, obtaining their speed as a function of time.

#### Noise and Dimensionality

To isolate the principal components (PCs) that capture significant fluctuations in neural activity, we first determined a ‘noisy’ firing rate for each neuron (*33, 37, 39*). This was achieved by calculating the firing rate of each neuron at time *t* during a trial within the condition *c*, and subtracting the average firing rate of all trials at the same time point *t* for that condition *c*. This process removed the signal component relevant to the neuron’s encoding function across conditions *c* and time, leaving only the residual fluctuations. Then, for each neuron, we randomly selected ten trials to compute the covariance with every other neuron. This random selection was repeated in each iteration to ensure a robust estimation of each neuron’s contribution to the overall noise covariance matrix, quantifying the covariance across all possible pairs of neurons. In mathematical notation, this follows:

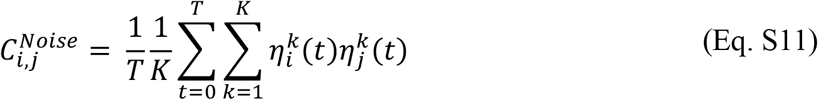

where 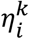 is the noisy firing rate value in time *t* of trial *k* that belongs to neuron *i*. PCA was then performed on the resulting covariance matrix to extract the eigenvalues, which indicate the variance captured by the principal components (PCs). We calculated the proportion of explained variance for each PC derived. To ensure statistical robustness, this matrix construction and eigenvalue extraction process was repeated 100 times with varying random seeds, allowing us to determine the mean and standard deviation of the noise-associated explained variance for the different PCs. This analysis enabled the identification of the dimensionality of the system through the identification of the meaningful PCs that had eigenvalues above the noise level, which were statistically significant. The significant PCs were then used to compute the corresponding projections.

#### Demixed Principal Component Analysis (dPCA)

Theoretical motivation and technicalities of this method are clearly explained in (*29, 32*). dPCA is divided into two steps: supervised and unsupervised. The supervised component of dPCA involves untangling the neural activity from pre-defined task variables in order to build marginalized covariance matrices, similar to variable selection in linear regression. Afterward, the method proceeds with an unsupervised principal component analysis on these matrices, similar to traditional PCA. In our task, we marginalized the population activity (*X*) across all liminal and supraliminal conditions (*X*_t,C_) for the tactile (5) and acoustic (5) modalities, excluding the absent stimulus condition. In this part of the study, we chose not to include absent stimuli when calculating the axes, allowing us to better understand the population dynamics between the two modalities, tactile and acoustic. However, even though the axes are computed without these stimuli, it remains possible to project the activity of the absent stimuli onto these axes (Figure 6).

To compute the demixed axes, the condition-independent component was subtracted during the marginalization process. To calculate the marginalization averages, we used the N-dimensional population activity:

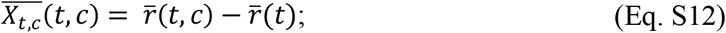

Then, 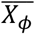 denotes the marginalized matrices with *ϕ* ∈ {{*t, c*} (the marginalization variable used in this study). Further, 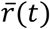 represents the firing rate average over all hit trials at each time bin (combining the ten conditions, 5 tactile and 5 acoustic), while 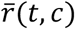 is the firing rate average per condition (separating trials according to the 5 different conditions within each sensory modality: tactile or acoustic). Once the marginalization is performed, dPCA finds separate decoder and encoder matrices for each 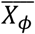 by minimizing the reduced-rank regression term:

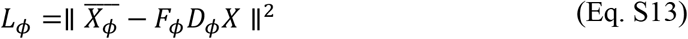

where 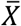 is the centered whole population data matrix (i.e., the average activity of each neuron is 0). The solution to this problem can be obtained analytically through singular value decomposition (SVD). Each component *D*_c_ can be ordered by the amount of explained variance, with the most prominent decoding axis referred to as the 1st demixed principal component (1st dPC or dPC1) of the variable *ϕ*. To avoid overfitting in dPCA, we introduced a regularization term and performed cross-validation to select the optimal regularization parameter. For Fig. 6A, we projected the N-dimensional data for a given condition 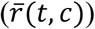 — in this case, the firing rate average per condition, computed by grouping trials according to tactile amplitude, acoustic amplitude, and stimulus-absent modalities — onto the most prominent decoding axis (*k*) or the variable *ϕ*.

Here we emphasize that dPCA finds the decoder and encoder of each marginalized variable, *ϕ*, minimizing each *L*_c_ separately. This allows for the independent calculation of the demixed decoder for the parameter condition. In Fig. 6A, we show the projections of neural activity sorted by condition onto the most prominent condition-dPCs. Most trajectories displayed an amodal encoding of tactile and acoustic stimuli, except along the condition-dPC4 & 5 axes, where the resultant representations exhibited a modality-specific (modal) coding of conditions across tactile and acoustic stimuli. Given that these modal axes show a clear separation between tactile and acoustic stimuli, we refer to them in Figure 6 as modal-dPCs. Conversely, the components that are independent of the modality are referred to as amodal-dPCs. In Fig. 6C, neural activity was averaged by stimulus modality (tactile or acoustic) or absence of stimulus (stimulus absent) and then projected onto the modal-dPC4 & 5. Unlike PCA, modal dPCA weights had a positive correlation (Fig. S4E). The projections were plotted against each other to create a 2D diagram, visualizing the separation of neural trajectories across modalities.

#### Regression Analysis with Multilinear Regression (MLR) and Generalized Linear Model (GLM)

We employed MLR (*41, 43*) and GLM to fit the mean firing rates of neuronal responses from different classes within the same sensory modality (tactile or acoustic) across various time windows. This analysis allowed us to assess the dynamic responses of neurons during specific periods of interest within the BDT.

The MLR model assumes that the mean firing rates are normally distributed. In our model, the regressors included tactile amplitude and acoustic amplitude, and the model was formulated as follows:

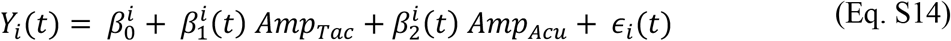

where *Y*_*i*_ (*t*) represents the mean firing rate of neuron *i* at time *t, Amp*_*TaC*_ (tactile amplitude) and *Amp*_*ACu*_ (acoustic amplitude) are the predictors, 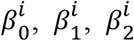 are the regression coefficients, *ϵ*_*i*_ (*t*) is the error term, assumed to be normally distributed. For the GLM, we assumed a Poisson distribution for the firing rates, with the identity link function. The GLM included additional regressors to account for the presence of tactile and acoustic stimuli which were specified as:

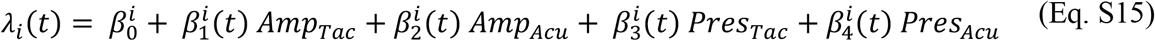

where *λ*_*i*_ (*t*) is the expected firing rate, *Pres*_*TaC*_ and *Pres*_*ACu*_ are binary indicators (0 or 1), representing whether any tactile or acoustic stimulus was present, respectively. Projection axes were constructed using the regression coefficients obtained from both MLR and GLM models. For each regression variable, a vector *r*_*m*_ was calculated and normalized such that ∥ *r*_*m*_ ∥ = 1. The mean firing rate for each condition, modality, and time bin was then projected onto these vectors using the dot product. In Fig. 5A, B, we employed GLM, while in Fig. S3C we employed MLR.

#### Recurrent Neural Network

We employed a recurrent neural network (RNN) (*30, 31, 41, 43, 47, 48, 50*) model to analyze neural response dynamics during BDT. The network architecture was inspired by previous studies, using an RNN described by the following dynamic equation:

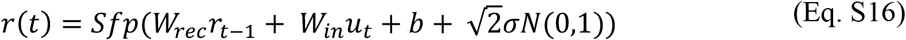

where *Sfp*(*x*) = *log* (1 + *e*^*x*^) is the softplus activation function, ensuring the non-negativity of the firing rates, *r*(*t*).

#### Network configuration

The network consisted of 256 units, each integrating signals through two primary pathways: Recurrent Weights (*W*_*reC*_), a matrix handling the recurrent connections, initialized with ones along the diagonal and other elements drawn from a normal distribution 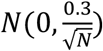, promoting stable dynamics (*31*); Input Weights (*W*_*in*_), a matrix for input connections that channels externals signals into the network; Bias and Noise, each unit incorporated a trainable bias *b* and Gaussian noise *N*(0,1) at each timestep to model variability and stochasticity in neuronal firing.

#### Input and stimulation

The input *u*_*t*_ to the RNN was structured to simulate sensory stimuli along with a go cue. Specifically, we defined three input units (as stimulus channels), one for each type of stimulus and one for the go cue. From 1 to 3.2 s post-trial start, one stimulus channel is switched on to one of the four amplitudes [0.01, 0.05, 0.1, 0.5], and is maintained for 0.5s. The go cue value remained at 0 until 1 second post-stimulus, whereupon it elevated to 1 until the trial’s end, half a second later.

#### Output layer and learning

The network activity was processed by a fully connected output layer *z*_*t*_, formulated as *z*_*t*_ = *W*_*out*_*r*_*t*_ + *bias*_*out*_, with each unit in this layer corresponding to a potential response (stimulus 1 present, stimulus 2 present, stimulus absent). The network was trained to minimize the squared error cost function *C* = ∑(*z*_*t*_ − *y*_*t*_)^2^ where *z*_*t*_ and *y*_*t*_ represents the activities of output units and target vectors, respectively. Parameters were updated via stochastic gradient descent, with a batch size of 64 and a learning rate of 0.0005. Training of the network proceeded unit performance accuracy, which we defined as correct answers produced (*30, 31*), reached 80%.

#### Rotating Memories

We propose modeling the sensory and memory retention dynamics observed in the VPC as a two-dimensional system of differential equations with state variables ***S*** and ***M***. These variables are intended to capture the dynamics along the two most representative modal dPCs—namely, dPC4 and dPC5 (Fig. 6). The first state variable, ***S***, models the stimulus-driven, early transient sensory response, while the second state variable, ***M***, represents the persistent mnemonic response, which retains the perceptual decision throughout the delay period.

To derive suitable dynamics for ***S*** and ***M***, we rely on the idea that sensory dynamics are predominantly faithful and linear, while mnemonic dynamics are categorical and nonlinear. Specifically, sensory responses exhibit exponentially vanishing memory, while mnemonic responses retain memory through multiple stable equilibria. Therefore, we propose that sensory dynamics can be modeled as a first-order low-pass filter driven by sensory inputs. In contrast, mnemonic dynamics are modeled as a first-order nonlinear system driven by the output of the sensory dynamics, featuring multiple stable equilibria: one corresponding to the VPC resting state (stimulus-absent response) and two others associated with the presence of acoustic and tactile stimuli.

The proposed model can be expressed as:

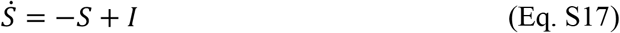

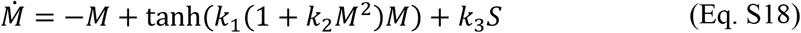

where *I* represents the sensory inputs (*I* < 0 for acoustic stimuli and *I* > 0 for tactile stimuli), and *k*_1_, *k*_2_, *k*_3_ are positive gain parameters. In line with our modeling hypothesis, the system described by these equations consists of two key components: a first-order filter with state variable ***S*** and a nonlinear multi-stable system with state variable ***M***. The low-pass filter in equation (Eq. S17) captures the transient nature of sensory responses, while the multi-stable nonlinear dynamics in equation (Eq. S18) model the categorical and persistent nature of mnemonic retention. The multi-stable structure of the mnemonic dynamics allows the system to maintain one of three stable states— stimulus-absent, tactile stimulus-present, or acoustic stimulus-present—thereby capturing the decision process observed in the VPC during the BDT (Fig. 8 & S5).

---

**Figure S1.**
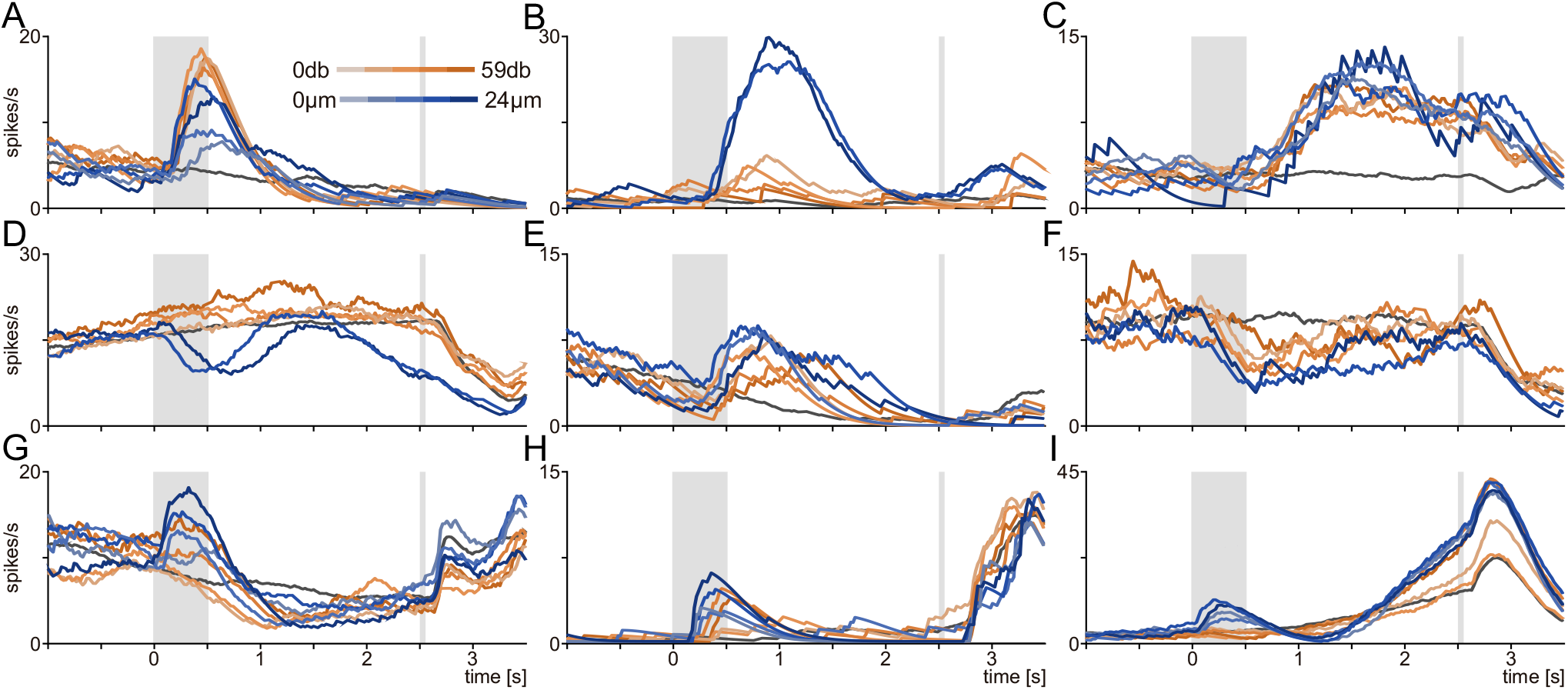
Showcase of the heterogeneous activity of 9 representative neurons (from A to I) recorded during BDT. Traces represent the firing rate average per class according to the amplitude of the sensory input presented (tactile and acoustic in tones of blue and orange, respectively) and for stimulus-absent trials (dark-gray traces). Light gray-shaded rectangles represent the stimulation period and PU event.

**Figure S2.**
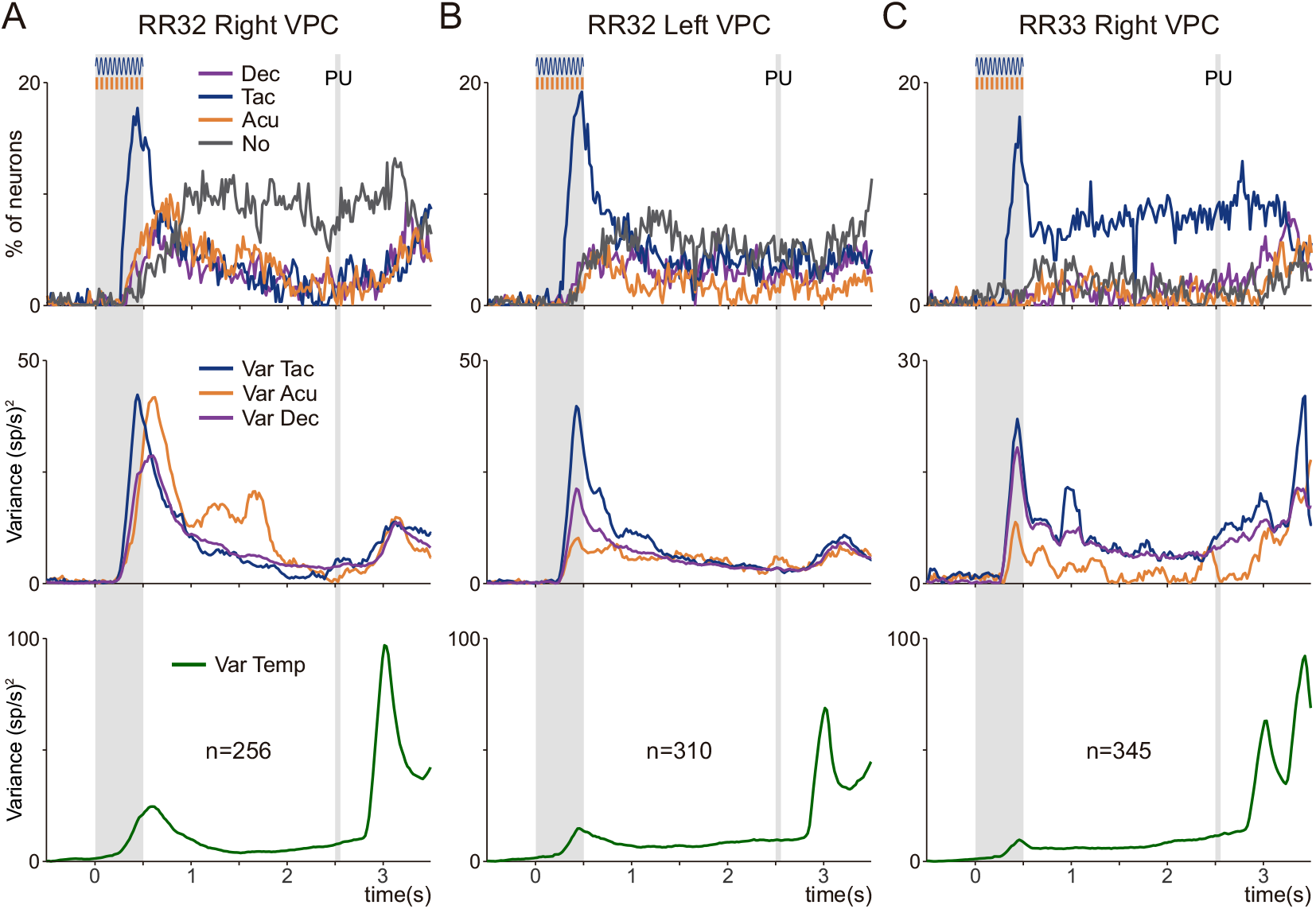
**(A-C)** Neuronal information processing types and cognitive and temporal variances across hemispheres: Right (n=256) and Left (n=310) in RR32, and Right (n=345) in RR33. Top: Proportion of neurons classified according to the matrix in Figure 3D, based on the significant information they convey about sensory input type—or its absence—and the corresponding decision. Consistent with previous observations, a majority of neurons in all hemispheres primarily encoded tactile information following stimulus onset. Middle: Variance in population responses over time, with blue traces for tactile (*Var*_*Tac*_) and orange for acoustic (*Var*_*Acu*_) stimulus classes, alongside purple traces for decision-making variance (*Var*_*Dec*_). Bottom: Temporal variances (*Var*_*Temp*_, green trace). Notably, post-PU, an increase in this metric is observed in the right hemispheres of both subjects, possibly linked to the ipsilateral orientation of this region to motor execution. In all panels, light gray-shaded rectangles denote the stimulation period and PU event.

**Figure S3.**
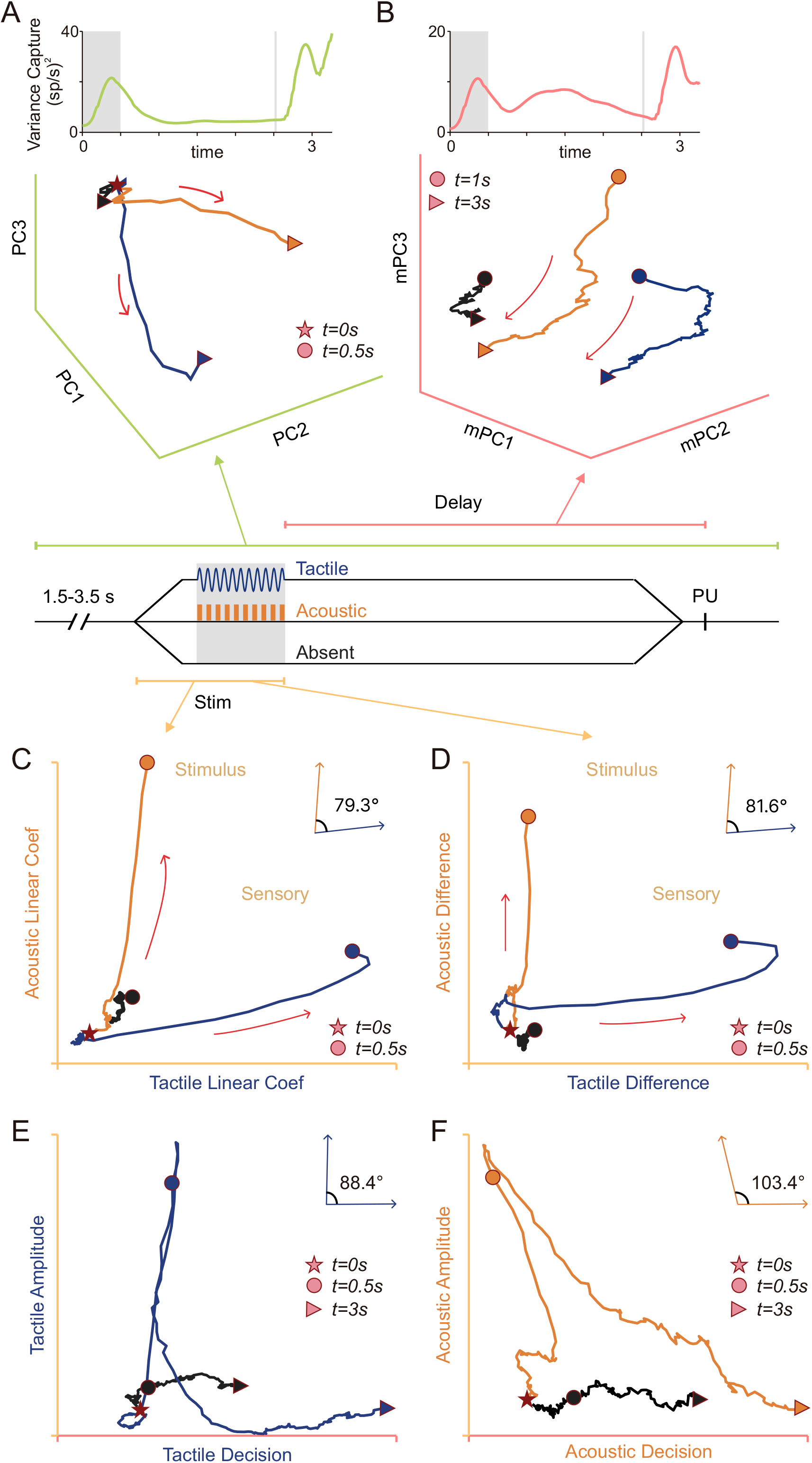
(A) Captured variance across time (top) and projected population dynamics (bottom), both using the first three PCs from PCA on the covariance matrix covering the entire BDT (−1 to 3.5 s, green line in BDT schematic). 3D diagrams were generated by projecting neuronal activity during the stimulation period (0 to 0.5 s, filled stars, and dots, respectively), sorted by modality (tactile [blue], acoustic [orange], or absent [black]). (B) Captured variance (top) and population dynamics (bottom) exhibited by the three mnemonic principal components (mPCs) obtained by applying PCA on the covariance matrix computed with the time-averaged modality-specific neural data in the delay period (0.5 to 2.5 s, pink line in task schematic). 3D diagrams were computed by projecting neural activity onto this mnemonic axis across delay and decision periods (1 to 3 s). (C) Projection of the decision averages (as in A and B) over axes obtained from an MLR, where the dependent variable was the mean firing rate of each class during the stimulus period, and the regressors were the tactile and acoustic amplitudes. Given that these regressors are not completely independent, the axes are not completely orthogonal (θ = 79.3°). (D) Similar to (C), but here the projection is over the normalized vectors obtained with the difference between the population vector pre-stimulus (t = 0) and post-stimulus (t = 0.5) (θ = 81.6). (E and F) Projections over decision and amplitude axes for the tactile (E) (θ = 88.4°) and acoustic (F) (θ = 103.4°) modalities. The amplitude axes were obtained as in panel C. The decision axes were obtained as in panel D but using the end of the delay period (t = 2.5) instead of the post-stimulus period.

**Figure S4.**
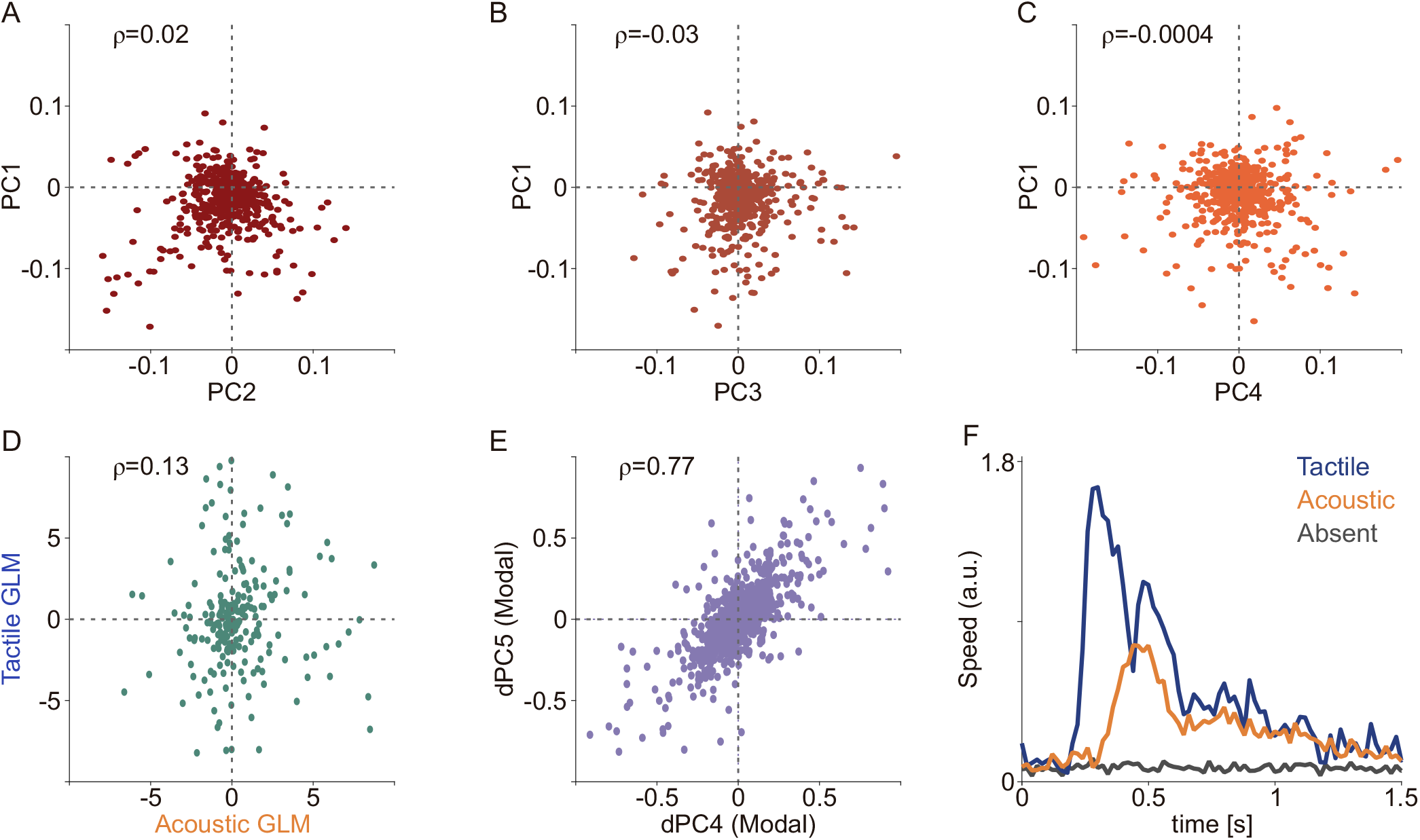
(A-E) Scatter plots of the weights associated with different neurons in some of the projections are shown. (A-C) Although the dynamics shown in the PCA in Figure 4 seem to be associated with different neuronal dynamics, no correlation is observed among the participating neurons. For example, the neurons involved in the tactile sensory dynamics (PCA1) appear to be uncorrelated with the neurons that separate the three dynamics during the delay (PCA4). (D) The sensory dynamics calculated with the GLM method (Fig. 5A) also show a low level of correlation among them. (E) The dynamics obtained with dPCA for modality, on the other hand, show a clear correlation. This coupling seems to be linked with the rotational dynamics observed in both the biological and simulated networks. (F) The speed of the population dynamics during the three types of stimuli is shown. Consistent with other results, the tactile dynamics respond earlier and faster. This suggests a greater capacity of the network to encode tactile stimuli.

**Figure S5.**
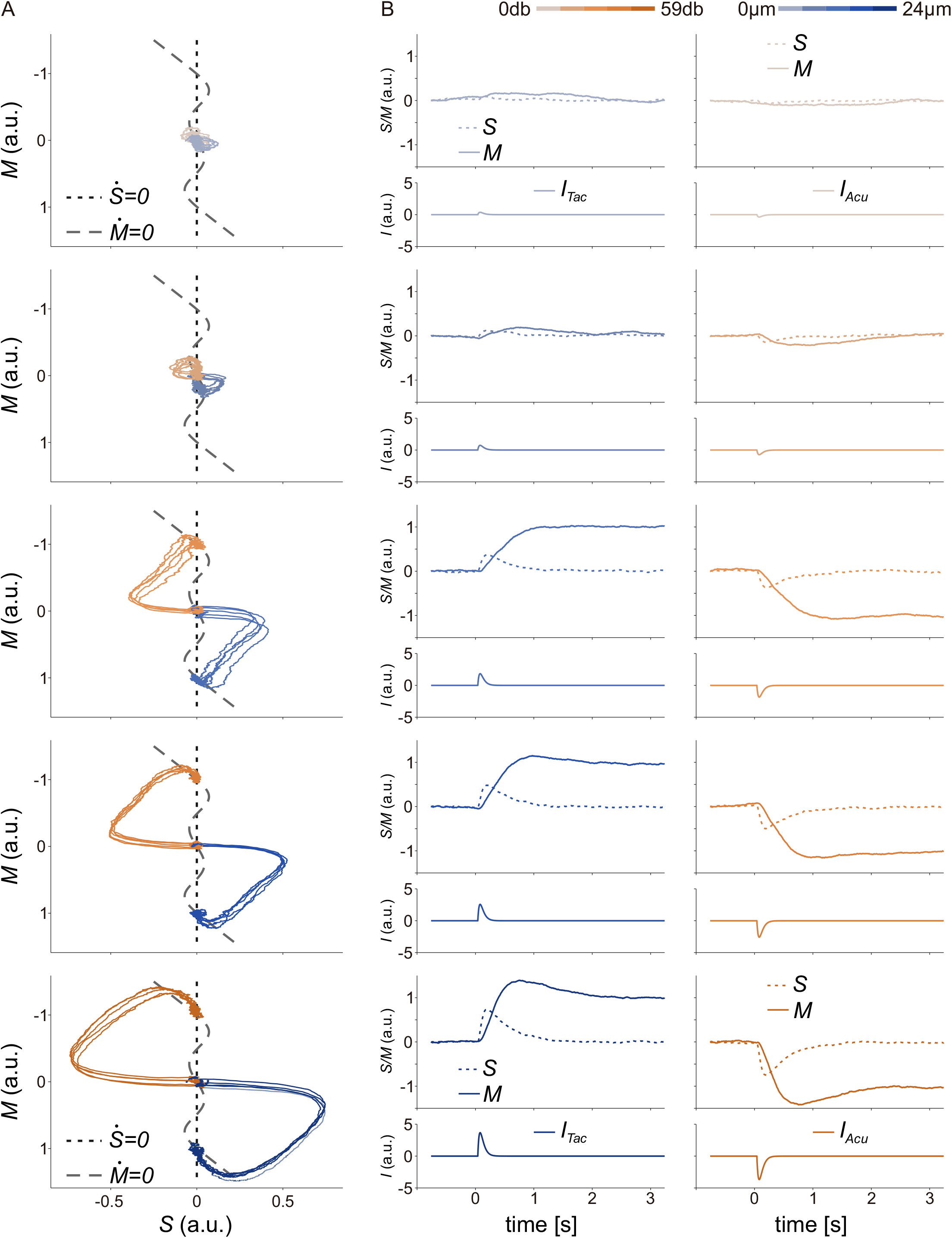
Responses of the low-dimensional model to stimuli of increasing amplitudes (A) Phase plane trajectories for tactile (blue) and acoustic (orange) stimuli of varying intensities. The S and M nullclines are indicated by dotted and dashed gray lines, respectively. (B) Time responses for both S (dotted line) and M (filled line) to the stimulus (I) are depicted at the bottom of each panel, with tactile trials on the left and acoustic trials on the right. For all stimulus amplitudes, the S variable returns to its initial state after the stimulation. Meanwhile, the M variable only returns when the stimulus is below the sensory threshold but keeps the memory of the presence and modality of the stimulus when the stimulus has a strong enough amplitude.

